# Divergence of olfactory receptors associated with the evolution of assortative mating and reproductive isolation in mice

**DOI:** 10.1101/2022.07.21.500634

**Authors:** Carole M. Smadja, Etienne Loire, Pierre Caminade, Dany Severac, Mathieu Gautier, Guila Ganem

## Abstract

Deciphering the genetic bases of behavioural traits is essential to understanding how they evolve and contribute to adaptation and biological diversification, but it remains a substantial challenge, especially for behavioural traits with polygenic architectures. In this study, we developed a population genomics approach coupled with functional predictions to address the evolution and genetic basis of olfactory-based assortative mate preferences in the house mouse, suspected to have evolved as a response to selection against hybridisation. We used whole genome resequencing data and the *C2* statistic of the program BayPass, which contrasts allele frequencies corrected for population structure, to characterize genetic differentiation between sets of populations with strong contrast in behaviour (expressing or not assortative mate preferences) and we identified some regions of the genome showing the expected significant and consistent association with behavioural divergence. A series of Olfactory and Vomeronasal Receptor genes, among the most differentiated genomic regions and in line with functional predictions, stand out as the prime candidates underlying this olfactory-based behavioural divergence. These genes form large gene clusters in the genome, with two main candidate clusters extending up to 1.8 Mb. Variant analyses indicate a potential dual role of regulatory and protein-coding changes in the evolution of choosiness. This study shows that combining expectations on the genomic patterns of divergence with functional expectations represents a promising route to unravelling the genetic architecture of complex trait variation and provides novel insights into the role of olfactory and vomeronasal receptors in mammal adaptation and speciation.

## Introduction

The evolution of behaviours plays a fundamental role in biological diversification, adaptation and speciation, but it remains a central challenge in biology to understand how behavioural traits evolve and the genetic basis of behavioural differences (Arguello and Benton 2017; Niepoth and Bendesky 2020; Jourjine and Hoekstra 2021). Behaviours involved in mate choice play a key role during the evolution of new species, as they act as premating components of reproductive isolation and may diverge rapidly through the action of direct and indirect selective effects (natural selection, sexual selection, reinforcing selection ( Servedio and Noor 2003; Smadja and Butlin 2011; Servedio and Boughman 2017; Kopp et al. 2018; Mendelson and Safran 2021). Some progress has been made, often by using quantitative genetics approaches, in characterising genes underlying assortative mate preferences in some biological systems (e.g. in *Drosophila* flies: Laturney & Moehring 2012, Pardy et al. 2021; sticklebacks: Bay et al. 2017; *Heliconius* butterflies: Merrill et al. 2019; European corn borer moths: Unbehend et al. 2021), but they represent cases where the genetic basis of behavioural divergence is relatively simple. Determining the genetic basis of behavioural divergence for more polygenic traits and identifying the signature of divergent selection acting on these underlying genomic regions remain more challenging.

In this study, we developed a population genomics approach based on the analysis of genetic differentiation between sets of populations with contrasted behaviours and selective pressures to address the evolution and genetic basis of olfactory-based assortative mate preferences in the house mouse. Population genomics has been successful in identifying targets of selection in the genome (Hohenlohe et al. 2010; Haasl and Payseur 2016) but its application to characterise the genetics of behavioural divergence remains largely unexplored (Kent et al. 2019; Niepoth and Bendesky 2020). With information on phenotype divergence among natural populations, on the proximal basis of the behavioural response, and on populations where these behaviours should be under selection, the house mouse system provides a predictive framework to address the evolution and genetics of behavioural divergence involved in reproductive isolation.

The two European subspecies of the house mouse (*Mus musculus musculus* and *M. m. domesticus*) diverged in allopatry ~350-500 kya from Central Asia/Middle East regions before coming into secondary contact in Europe ~5 kya (Boursot et al. 1996; Geraldes et al. 2011). Some degree of postzygotic isolation has been demonstrated in this hybrid zone, mostly in the form of reduced hybrid fertility (Britton-Davidan et al. 2005; Albrechtova et al. 2012; Turner et al. 2012). However, the two subspecies also display some degree of sexual isolation where they meet and form a hybrid zone: assortative mate preferences (accompanied by a propensity to mate with the preferred partner) have been evidenced in both sexes of both subspecies, though preference is stronger in *M. m. musculus* (Smadja and Ganem 2002; Smadja et al. 2004; Smadja and Ganem 2005; Ganem et al. 2008; Bimova et al. 2011; Latour et al. 2014; Smadja et al. 2015).

Several arguments suggest that these assortative mate preferences have evolved under selection, as a result of a reinforcement process (Servedio and Noor 2003; Butlin and Smadja 2018). Within each subspecies, assortative mate preferences are only displayed in populations from the hybrid zone (‘Choosy’ populations) where selection against hybridization is known to act, while populations from allopatry do not display any significant directional preferences (‘Non-Choosy’ populations) (Smadja and Ganem 2005; Smadja and Ganem 2008; Bimova et al. 2011; Latour et al. 2014; Smadja et al. 2015). This pattern of reproductive character displacement suggests a mechanism of reinforcement, by which assortative mate preferences would have evolved as an adaptive response to the existing postzygotic barriers in this hybrid zone. In this study, we used this information to identify the genes underlying the evolution of assortative mate preferences by articulating and testing predictions on the patterns of genomic divergence expected for loci experiencing reinforcing selection. One key prediction is that genetic divergence will be significantly elevated at loci underlying assortative mate preferences when sympatric (Choosy) and allopatric (Non-Choosy) individuals of the same species are compared (Garner et al. 2018).

To identify loci showing this expected pattern of divergence, we leveraged the high-quality reference genome available for this species (Waterston et al. 2002), produced whole-genome resequencing data for the same individuals and populations that had been behaviourally typed previously (Smadja et al. 2015), and finely characterised genomic divergence between Choosy and Non-Choosy populations within *M. m. musculus*, the subspecies where the contrast in behaviour is the strongest between these two types of populations (Smadja et al. 2004; Smadja and Ganem 2005; Ganem et al. 2008; Latour et al. 2014). We opted for a pool-sequencing strategy as a cost-effective way to assess genetic differentiation among populations with contrasted behaviours (Bastide et al. 2013; Yang et al. 2015), particularly relevant in our case where mate preferences are statistically assessed in wild mice at the population level and show limited variability among individuals within populations (Smadja et al. 2004; Smadja and Ganem 2005; Smadja et al. 2015). To ensure the identification of genetic variants specifically divergent between Choosy and Non-Choosy populations, we combined multiple population comparisons (focal and control comparisons between groups of populations with or without contrasted behaviours) and a replicated design within each group compared. We used the program BayPass and its recently implemented *C2* contrast statistic to identify outlier loci under selection between Choosy and Non-Choosy populations and associated with the behavioural contrast (Gautier 2015; Olazcuaga et al. 2021) (Figure 1 and Methods). The Bayesian hierarchical model implemented in BayPass represents a flexible and powerful framework to carry out association analyses and scans for selection since it efficiently accounts for the neutral correlation structure among allele frequencies in the sampled populations. In addition, it allows handling properly Pool-Seq data and accounts for the additional level of variation they introduce in the estimation of the parameter of interest. .

**Figure 1:**
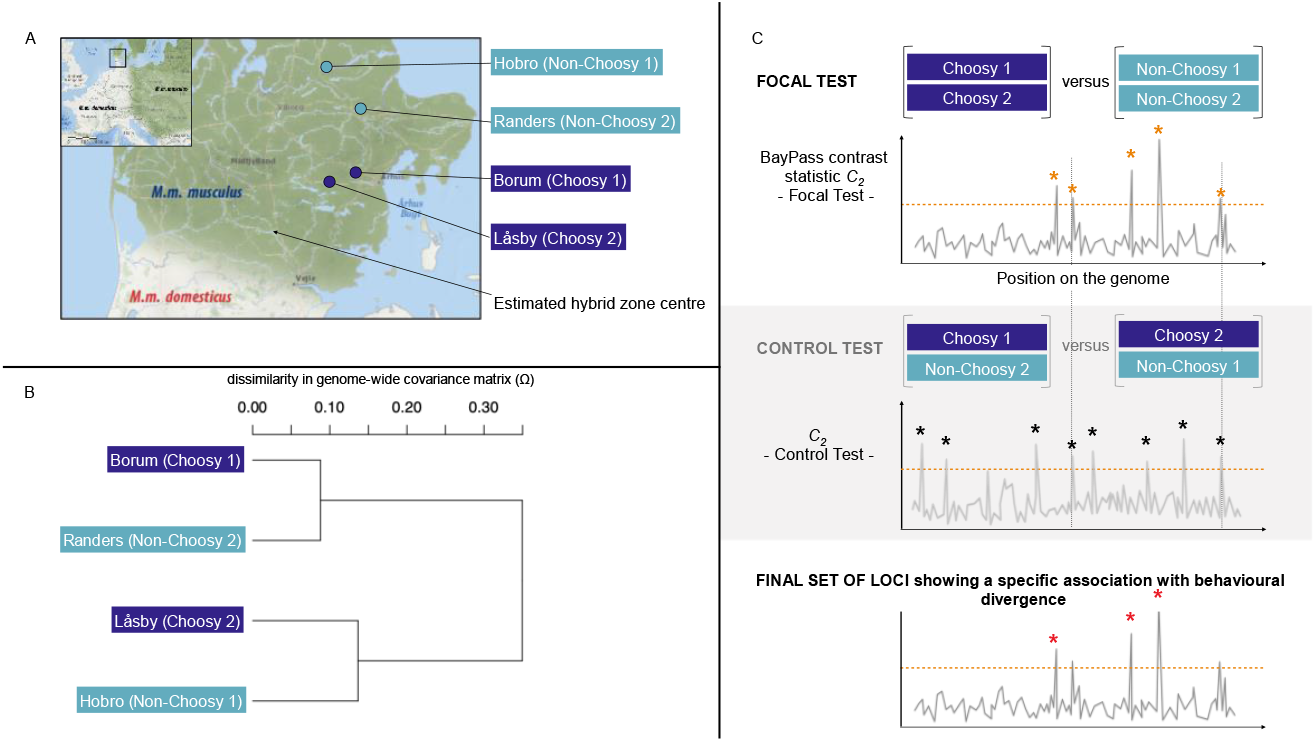
Sampling and experimental design. A) Populations of Mus musculus musculus sampled in localities in the M. m. musculus - domesticus hybrid zone (dark blue points) or further away from the centre of the hybrid zone (light blue points). Mean nucleotide diversity: Låsby (n=27, π = 0.002 +/- 1.136e-05), Borum (n=33, π = 0.003 +/- 1.151e-05), Randers (n=29, π = 0.002 +/- 1.027e-05), Hobro (n=29, π = 0.002 +/- 1.159e-05); pairwise FST estimates between Choosy and Non-Choosy populations: FST Borum-Choosy1 - Randers-NonChoosy2 = 0.057; FST Låsby-Choosy2 - Randers-NonChoosy2 = 0.087; FST Hobro-NonChoosy1 - Borum-Choosy1 = 0.089; FST Hobro-NonChoosy1 - Låsby-Choosy2 = 0.114); FST within behavioural groups: FST Borum-Choosy1 - Låsby-Choosy2 = 0.065; FST Hobro-NonChoosy1 - Randers-NonChoosy2 = 0.076. B) Relatedness among populations based on the genome-wide covariance matrix (Ω) across population allele frequencies for autosomal chromosomes (similar results for the X chromosome). C) The two series of population comparisons were used to identify loci showing an association with behavioural divergence. The Focal Test compared replicated sets of Choosy and Non-Choosy populations (Choosy1+Choosy2 versus Non-Choosy1+Non-Choosy2) to identify loci showing significant divergence (as measured by the contrast statistic C2 from the BayPass program) between populations with contrasted behaviour. The Control Test compared two other combinations of the same populations, this time not grouped by their behavioural profile (Choosy1+Non-Choosy2 versus Choosy2+Non-Choosy1). Only loci found divergent specifically in the Focal Test were retained as the final list of candidate loci.

We applied this combination of population comparisons for two types of outlier analyses at different genomic scales: SNP-based differentiation across the whole genome (based on the estimation of SNP-based *C_2_* contrast statistic), and gene-based differentiation across all annotated genes. While most existing statistical methods for detecting outliers focus on analysing SNPs one-by-one, SNP-based analyses may have some limitations, among which a loss of power due to stringent correction applied to limit false positives in this large number of comparisons (millions of SNPs in a genome) and weak sensitivity to detecting multiple causal variants effects. Leveraging methods developed in the context of GWAS in humans (Neale and Sham 2004; Huang et al. 2011; Chung et al. 2019), we complemented SNP-based analyses with gene-based analyses to gain power in detecting outliers by aggregating disparate signals within a gene as well as to make more direct link of the outlier analyses with the phenotype of interest. To capture different potential underlying genetic architectures of adaptive divergence (single causal variant versus multiple weakly selected variants within a gene), we analysed two commonly used gene-based statistics representing the highest SNP-based differentiation value of a gene and the average differentiation value of a gene (Wang et al. 2017; Dickson et al. 2020), called respectively *C_2_max_* and *C_2_mean_* in our study.

Not all differentiated genomic regions between Choosy and Non-Choosy populations are expected to be associated with the behavioural trait of interest, since other traits and factors can differ between this set of populations. To further interpret genomic patterns of divergence and link phenotypic and genomic variation, we therefore combined this population genomics approach with functional expectations (Haasl and Payseur 2015; Ravinet et al. 2017; Wolf and Ellegren 2017), by leveraging existing knowledge on the implication of olfaction in the expression of these assortative mate preferences in the house mouse. Previous behavioural assays and chemical analyses have shown that subspecies recognition and associated assortative mate preferences could be driven (at least partly) by olfactory cues present in the mouse urine (Smadja and Ganem 2008; Hurst et al. 2017) and that urinary cues are sufficient to elicit mate choice between the two subspecies (Smadja and Ganem 2002; Smadja et al. 2004; Smadja and Ganem 2005; Ganem et al. 2008; Bimova et al. 2011). The olfactory system thus plays a major role in the expression of assortative mate choice between the two subspecies, and we expect some genes involved in odorant recognition to be associated with the evolution of sexual isolation in the house mouse hybrid zone. This includes olfactory receptor (OR) genes expressed in the main olfactory epithelium and primarily involved in the recognition of airborne chemicals present in the environment (odorants and pheromones), and vomeronasal receptor (VR) genes expressed in the vomeronasal organ and known to play a key role in both volatile and non-volatile kairomone and pheromone recognition (Isogai et al. 2011; Chamero et al. 2012; Hayden and Teeling 2014; Liberles 2014; Bear et al. 2016). These genes are organized in very large multi-gene families in the mouse genome (more than 1,200 ORs and 530 VRs (Zhang and Firestein 2002; Ibarra-Soria, M. Levitin, et al. 2014; Ibarra-Soria, M.O. Levitin, et al. 2014)). Here, we considered those genes as key candidates among the possible genetic determinants of assortative mate preferences and used this list of candidate genes to interrogate outlier regions and narrow down regions of interest.

Patterns of genomic differentiation between Choosy and Non-Choosy populations confirmed that some regions of the genome show the patterns of divergence expected under selection. Among those, a handful of olfactory and vomeronasal receptors showed genetic divergence associated with the observed behavioural divergence between hybrid zone and allopatric populations. Our study thus provides new insights into the role of olfactory and vomeronasal receptors in mammal adaptation and speciation. It also offers a novel experimental framework to identify loci showing the expected patterns of divergence under reinforcement, and confirms that combining expectations on the genomic patterns of divergence with functional expectations is a promising route to decipher the genetic architecture of behavioural trait variation.

## Results

All supplementary figures and tables are available online (Smadja et al. 2022a).

### Sequence divergence between Choosy and Non-Choosy populations

Whole-genome resequencing of pools of individuals representing two Choosy and two Non-Choosy populations of *M. m. musculus* (Table S2, and see Figure 1a and Table S1 for a detailed sample description) identified a total of 28,500,265 SNPs in the dataset after stringent filtering. Genetic dissimilarity indexes estimated from the scaled genome-wide covariance matrix of the population allele frequencies (Ω matrix, produced by the program BayPass) indicated that the two Choosy populations do not cluster together (Figure 1b). This result could be explained by the dense river network in Jutland acting as geographic barriers to migration and hence influencing population structure (Raufaste et al. 2005). Interestingly, this pattern of population relatedness makes the identification of outlier divergent loci in the Focal Test comparing Choosy and Non-Choosy populations (Figure 1c) conservative. A Control Test was performed to remove outlier loci from the Focal Test representing population-specific divergence and therefore not a consistent divergence between both Choosy versus both Non-Choosy populations (Figure 1c and Methods).

#### Outlier SNPs

We confirmed, in both Focal and Control Tests, that the SNP-based *C_2_* contrast statistic follows a *χ*^2^ distribution with one degree of freedom as expected under the null hypothesis (Figure S1). We thus calculated *χ*^2^-associated *p*-values and FDR to identify outlier SNPs showing extreme genetic differentiation in each population comparison (Focal and Control). We identified 1,612,787 outlier SNPs (*P* < 0.05) in the Focal Test before multiple testing corrections, reduced to 6546 outlier SNPs after applying FDR correction. In the Control Test, 1,111,516 SNPs were identified as outliers before multiple testing corrections. We retained as the final list of ‘behavioural’ outlier SNPs those 4818 SNPs showing both significant differentiation between Choosy and Non-Choosy populations after FDR correction in the Focal Test and no significant differentiation in the Control Test (Table S3).

#### Outlier genes

In the Focal Test, at the *P* < 0.05 threshold, we identified 999 *C_2_max_* outlier genes (over 41,702 annotated genes included in this permutation test) (Table S4a) and 1756 *C_2_mean_* outlier genes (over 39,598 annotated genes included in this analysis) (Table S4b). In the Control Test, comparable numbers of outlier genes were found: 1108 *C_2_max_* outlier genes (Table S4c) and 1612 *C_2_mean_* outlier genes (over 39,447 tested genes, among which 1472 genes with no permutation results due to insufficient SNP number or too large gene size) (Table S4d). However, the large majority of genes found significant in the Focal Test were not found significant in the Control Test: 77 genes had significant *C_2_max_* in both the Focal and the Control Tests (considered as non-consistently associated with the behavioural contrast); as a result, from the 999 *C_2_max_* initial outlier genes in the Focal Test, 922 were retained as the final *C_2_max_* outlier genes; similarly, out of the 1756 initial *C_2___mean_* outlier genes, 1622 were retained as the final *C_2_mean_* outlier genes after removing 134 outliers not consistently associated with the behavioural contrast. The union of these two cured outlier gene sets, composing the final and complete list of outlier genes, included 2118 genes, among which 496 were *C_2_max_* specific outliers, 1196 were *C_2_mean_* specific outliers and 426 were *C_2_max_* and *C_2_mean_* outlier genes (Table S5).

Among those 2118 outlier genes, we categorised the most strongly differentiated ones between Choosy and Non-Choosy populations according to (1) their level of differentiation at one of the two gene-based contrast statistics (*p*-value *C_2_max_* < 0.01 or *p*-value *C_2_mean_* < 0.01), hereafter called highly significant outlier genes or (2) their level of differentiation at both gene-based statistics (see Methods and Table S5 for details), hereafter called the ‘top’ outlier genes. We found a total of 479 highly significant outlier genes, among which 111 *C_2_max_* specific outliers, 142 *C_2_mean_* specific outliers and 226 *C2_max+C2_mean* outlier genes (Table S5, bold gene_IDs). Among these highly significant outlier genes, we further identified 95 outlier genes showing consistent signals of strong differentiation at both *C_2_max_* and *C_2_mean_* gene-based contrast statistics, that were consequently classified as ‘top outlier genes’ (Table S5, bold and underlined gene_IDs).

### Functional impact and nature of candidate genomic regions associated with behavioural divergence

#### Genic and regulatory regions primarily associated with behavioural divergence

Using the Ensembl Variant Effect Predictor (VEP), we found that the 4818 outlier SNPs identified as significantly differentiated between Choosy and Non-Choosy populations include 1390 novel variants and 3428 already annotated variants in the Ensembl database (71.1%). Those outlier SNPs overlap 977 annotated genes, 2878 transcripts and 412 regulatory features (Table S6). Genic variants represent a majority of those outlier SNPs (54.03%), with 50.76% of intronic variants and the rest corresponding to exonic (1.31%) and UTRs (1.96%) variants (Table S6, Figure 2a). Protein-coding variants include 64% of synonymous variants, 32% of missense variants and 3% of stop-gained variants (respectively 0.48%, 0.24% and 0.02% of the total, Figure 2a). Non-genic variants are slightly less represented (45.97%) but include a large number of variants (32.91% of all outlier SNPs) potentially involved in gene regulation: down- and upstream variants as well as already known regulatory region variants (Table S6, Figure 2a). Only 13% of all outlier SNPs fall in intergenic regions. In comparison, predictions made on all SNPs (and not only outlier SNPs) in the dataset show fewer variants potentially involved in gene regulation (24%), fewer UTR variants (0.70%), fewer synonymous and missense variants (0.37% and 0.19% respectively) and 10 times less stop-gained variants (0.0022%). In contrast, they show more intergenic and intronic variants (16.15% and 57.62%, respectively).

**Figure 2:**
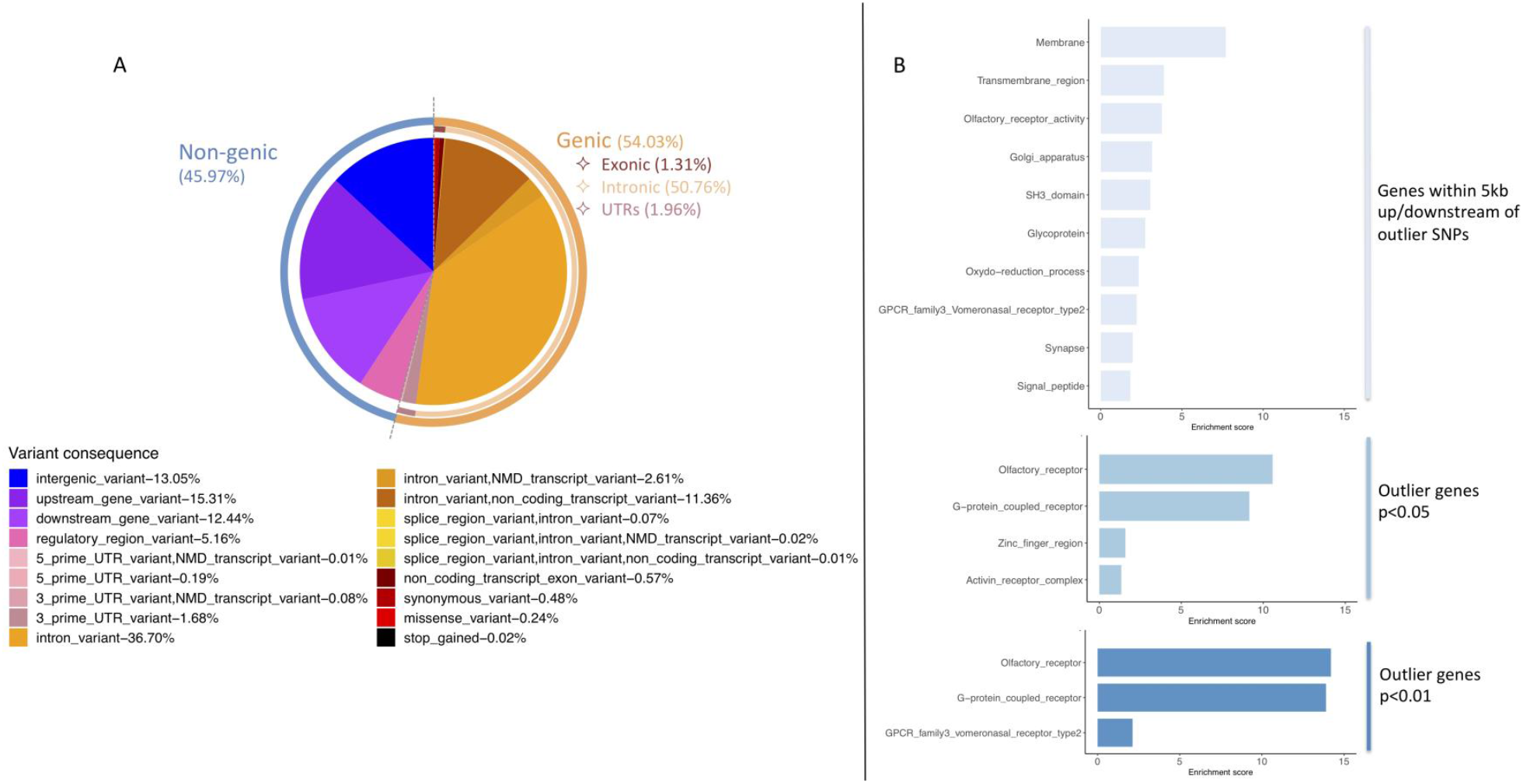
Functional analysis results. A) Summary of Ensembl VEP analysis performed on all outlier SNPs. The pie chart shows the proportions of the different variant consequences found in this set of outlier SNPs. Consequences include information on the location and possible functional effect of each variant (e.g. intergenic, within 5k up-/down-stream of an annotated genes, UTRs, intronic, exonic, synonymous, missense, stop_gained, as well as known regulatory regions). B) DAVID functional enrichment analyses. Up to top 10 functional annotation clusters (enrichment score > 1.3) are shown for analyses performed on (i) genes found within 5 kb of outlier SNPs (ii) genes identified as outlier genes (C2_max and/or C2_mean) at 0.05 threshold, (iii) genes identified as outlier genes at 0.01 threshold.

#### Olfactory and vomeronasal receptor genes as prime candidates

Functional enrichment analyses using the program David were performed on four gene datasets: genes within 5 kb of outlier SNPs (977 initial genes, 656 *Ensembl* gene IDs passing David database criteria), the complete list of outlier genes (2118 initial genes, 1181 David gene IDs), the list of highly significant outlier genes (479 initial genes, 291 David gene IDs) and the list of ‘top’ outlier genes (95 initial genes, 64 David gene IDs) (see Table S5 for a complete list of outlier genes, with their gene names and gene IDs). For the gene set within 5 kb of outlier SNPs, we found 14 highly significant functional annotation terms (Table S7a) and 17 significant functional annotation clusters (Table S7b). The three most enriched functional annotation clusters were membrane proteins, trans-membrane region proteins and olfactory receptor activity; Golgi apparatus, SH3 domain, glycoproteins, reduction-oxidation process and synapse were also part of enriched annotation clusters (Figure 2b). For outlier gene sets, olfactory receptor types of function were systematically found the most significantly enriched and enrichment score increased as the set of outlier genes is narrowed down to the most significant outlier genes (Figure 2b, Table S7c-h). For the gene sets corresponding to the most promising candidates, we found a very highly significant enrichment of only one group of functional categories corresponding to olfactory receptor functions (olfactory receptor/ G-protein-coupled receptor/ GPCR family3 vomeronasal receptor type2) (Figure 2b & Table S7e-f: highly significant outlier genes *p* < 0.01, Table S7g-h: ‘top’ outlier genes).

A total of 127 Olfactory Receptor (*Olfr*) genes and 27 Vomeronasal Receptor (*Vmn*) genes were identified as outlier genes (mostly on chromosomes 2, 7 and 9), and 55 *Olfr* and 13 *Vmn* are among the highly significant (*p* < 0.01) outlier genes (Table 1 and Table S5). Consistent with David functional enrichment results, this corresponds to an over-representation of *Olfr* genes among the complete list of outlier genes (hypergeometric test, *n* _Olf outliers_ = 127, *n* _Olf all_ = 1356; n _genes outliers_ = 2118; *n* _genes all_ = 50942, *p*-value = 1.90 e-17), and an over-representation of both *Olfr* genes (*n* _Olf outliers_ = 55, *n* _Olf all_ = 1356; n _genes outliers_ = 479; *n* _genes all_ = 50942, *p*-value = 1.37 e-19) and *Vmn* genes (*n* _vmn outliers_ = 13, *n* _vmn all_ = 543; n _genes outliers_ = 479; *n* _genes all_ = 50942, *p*-value = 0.0021) among highly significant outlier genes. Among Vomeronasal Receptor gene candidates, 10 belong to family 1 of Vomeronasal Receptors and 17 to family 2 of Vomeronasal Receptors (Yang and Zhang 2007; Zhang et al. 2007), the proportion of family 2 *Vmn* genes increasing when reducing the outlier gene set to the most significant ones (number of family 1 *Vmn* genes = 1, number of family 2 *Vmn* genes = 12) (Table 1 and Table S5). The majority of *Olfr* outlier genes are *C_2_mean_* outliers while the majority of *Vmn* outlier genes are *C_2_max_* outliers (Table S5). Among the ‘top’ outlier genes defined when combining information from the *C_2_mean_* and *C_2_max_* statistics, we found 23 *Olfr* and 4 *Vmn* genes (Table 1 and Table S5).

**Table 1:**
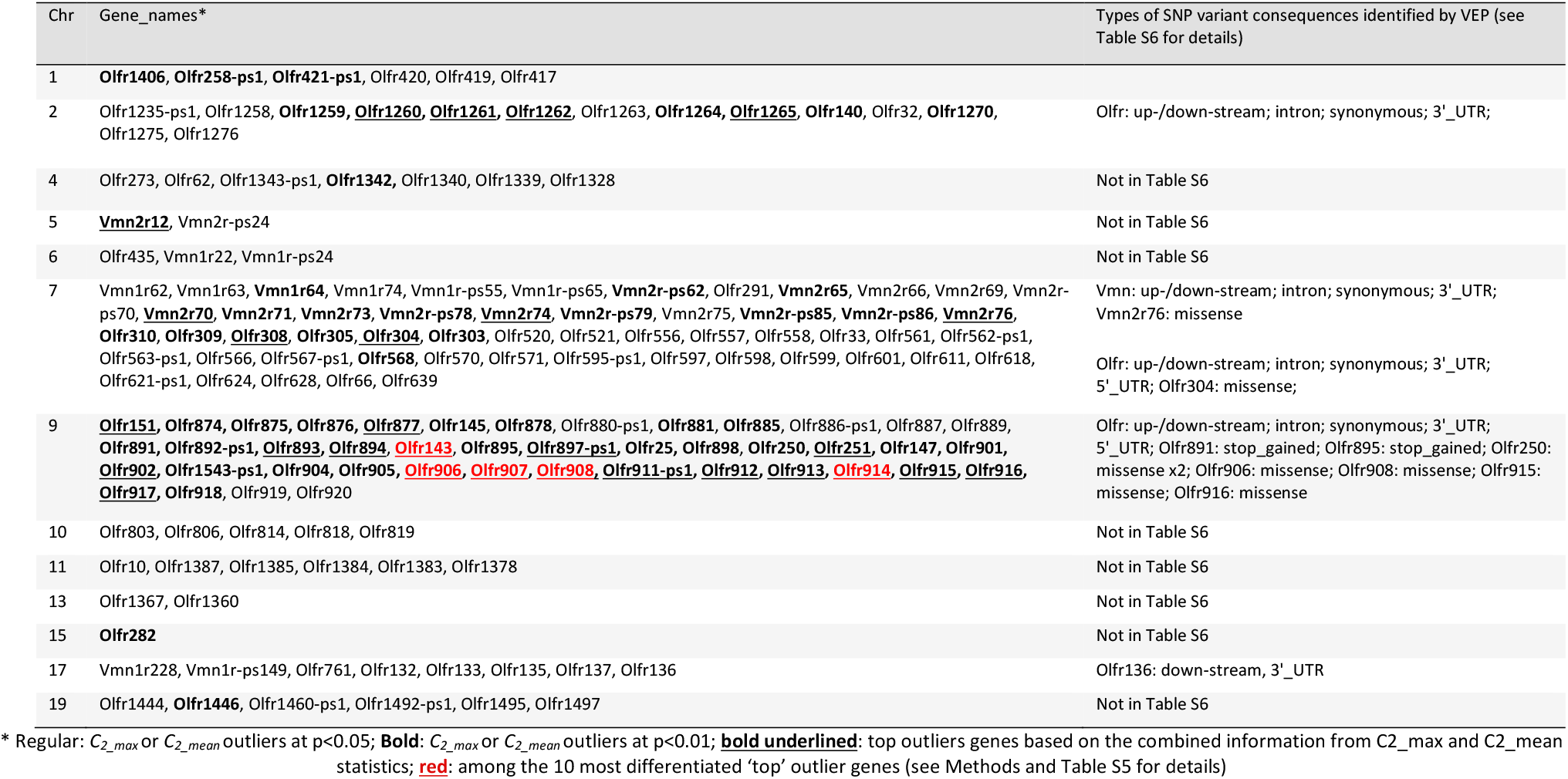
List of significantly differentiated Olfactory (Olfr) and Vomeronasal (Vmn) Receptor genes between Choosy and Non-Choosy populations

| Chr | Gene_names* | Types of SNP variant consequences identified by VEP (see Table S6 for details) |
| --- | --- | --- |
| 1 | **Olfr1406**, **Olfr258-ps1**, **Olfr421-ps1**, Olfr420, Olfr419, Olfr417 |  |
| 2 | Olfr1235-ps1, Olfr1258, **Olfr1259**, **Olfr1260**, **Olfr1261**, **Olfr1262**, Olfr1263, **Olfr1264**, **Olfr1265**, **Olfr140**, Olfr32, **Olfr1270**, Olfr1275, Olfr1276 | Olfr: up-/down-stream; intron; synonymous; 3' _UTR; |
| 4 | Olfr273, Olfr62, Olfr1343-ps1, **Olfr1342**, Olfr1340, Olfr1339, Olfr1328 | Not in Table S6 |
| 5 | **Vmn2r12**, Vmn2r-ps24 | Not in Table S6 |
| 6 | Olfr435, Vmn1r22, Vmn1r-ps24 | Not in Table S6 |
| 7 | Vmn1r62, Vmn1r63, **Vmn1r64**, Vmn1r74, Vmn1r-ps55, Vmn1r-ps65, **Vmn2r-ps62**, Olfr291, **Vmn2r65**, Vmn2r66, Vmn2r69, Vmn2r-ps70, **Vmn2r70**, **Vmn2r71**, **Vmn2r73**, **Vmn2r-ps78**, **Vmn2r74**, **Vmn2r-ps79**, Vmn2r75, **Vmn2r-ps85**, **Vmn2r-ps86**, **Vmn2r76**, **Olfr310**, **Olfr309**, **Olfr308**, **Olfr305**, **Olfr304**, **Olfr303**, Olfr520, Olfr521, Olfr556, Olfr557, Olfr558, Olfr33, Olfr561, Olfr562-ps1, Olfr563-ps1, Olfr566, Olfr567-ps1, **Olfr568**, Olfr570, Olfr571, Olfr595-ps1, Olfr597, Olfr598, Olfr599, Olfr601, Olfr611, Olfr618, Olfr621-ps1, Olfr624, Olfr628, Olfr66, Olfr639 | Vmn: up-/down-stream; intron; synonymous; 3' _UTR; Vmn2r76: missense<br>Olfr: up-/down-stream; intron; synonymous; 3' _UTR; 5' _UTR; Olfr304: missense; |
| 9 | **Olfr151**, **Olfr874**, **Olfr875**, **Olfr876**, **Olfr877**, **Olfr145**, **Olfr878**, Olfr880-ps1, **Olfr881**, **Olfr885**, Olfr886-ps1, Olfr887, Olfr889, **Olfr891**, **Olfr892-ps1**, **Olfr893**, **Olfr894**, **Olfr143**, **Olfr895**, **Olfr897-ps1**, **Olfr25**, **Olfr898**, **Olfr250**, **Olfr251**, **Olfr147**, **Olfr901**, **Olfr902**, **Olfr1543-ps1**, **Olfr904**, **Olfr905**, **Olfr906**, **Olfr907**, **Olfr908**, **Olfr911-ps1**, **Olfr912**, **Olfr913**, **Olfr914**, **Olfr915**, **Olfr916**, **Olfr917**, **Olfr918**, Olfr919, Olfr920 | Olfr: up-/down-stream; intron; synonymous; 3' _UTR; 5' _UTR; Olfr891: stop_gained; Olfr895: stop_gained; Olfr250: missense x2; Olfr906: missense; Olfr908: missense; Olfr915: missense; Olfr916: missense |
| 10 | Olfr803, Olfr806, Olfr814, Olfr818, Olfr819 | Not in Table S6 |
| 11 | Olfr10, Olfr1387, Olfr1385, Olfr1384, Olfr1383, Olfr1378 | Not in Table S6 |
| 13 | Olfr1367, Olfr1360 | Not in Table S6 |
| 15 | **Olfr282** | Not in Table S6 |
| 17 | Vmn1r228, Vmn1r-ps149, Olfr761, Olfr132, Olfr133, Olfr135, Olfr137, Olfr136 | Olfr136: down-stream, 3' _UTR |
| 19 | Olfr1444, **Olfr1446**, Olfr1460-ps1, Olfr1492-ps1, Olfr1495, Olfr1497 | Not in Table S6 |

\* Regular: $C_{2\_max}$ or $C_{2\_mean}$ outliers at $p < 0.05$ ; **Bold**: $C_{2\_max}$ or $C_{2\_mean}$ outliers at $p < 0.01$ ; **bold underlined**: top outliers genes based on the combined information from $C_{2\_max}$ and $C_{2\_mean}$ statistics; **red**: among the 10 most differentiated 'top' outlier genes (see Methods and Table S5 for details)

Beyond the over-representation of Olfactory Receptor and Vomeronasal Receptor genes among outlier gene sets, some of them appeared among the top most significantly differentiated outlier genes: 10 Olfr and Vmn are among the 30 most differentiated C2_max outlier genes (Olfr304, Olfr907, Olfr911-ps1, Olfr913, Olfr915, Olfr908, Olfr914, Olfr143, Olfr906 and Vmn2r74) and 8 Olfr among the 30 most differentiated C2_mean outlier genes (Olfr906, Olfr908, Olfr914, Olfr907, Olfr143, Olfr151, Olfr894 and Olfr1265) (Table S5). Five Olfr genes, all in a gene cluster on chromosome 9, were common to these two lists and among the 10 most differentiated ‘top’ outlier genes: Olfr906, Olfr907, Olfr908, Olfr914 and Olfr143, with Olfr906, Olfr907 and Olfr908 among the 4 most differentiated ‘top’ outliers (Table 1 and Table S5). Comparatively to Olfr genes, Vmn outlier genes were proportionally less represented among the most differentiated genes, but four of them were found in the list of ‘top’ outlier genes: Vmn2r74, Vmn2r76, Vmn2r70 and Vmn2r12 (Table S5).

Table 1 also reports, when available from VEP results (Table S6), the functional impact of outlier SNPs found in candidate outlier *Vmn* and *Olfr* genes. The large majority of reported functional consequences corresponded to up- and down-stream variants, UTR variants, intronic variants and synonymous variants. However, five *Olfr* genes on chromosome 9 (*Olfr906, Olfr908, Olfr915, Olfr916* and *Olfr250*) and two receptor genes on chromosome 7 (*Olfr304* and *Vmn2r76*) additionally showed a missense mutation between Choosy and Non-Choosy populations, and *Olfr891* and *Olfr895* on chromosome 9 showed a stop-gain variant.

Genes that are not Olfactory or Vomeronasal Receptor genes were also among outlier genes (Table S5). Most of them do not have a well-characterized function, and when they do, their link with the behavioural phenotype of interest is unclear. We nevertheless noted some genes involved in gene regulation (*Khsrp*, among the four most differentiated ‘top’ outlier genes), regulation of neural genes (*Barhl2*), acting as transcription factor (*Gtf2f1*), or expressed in testes and potentially inducing male infertility (*Tex21, Dazl*).

### Genetic characteristics of regions associated with behavioural divergence

#### Association with some other genomic features

*No association with regions of reduced genetic diversity and coldspots of recombination :* genetic diversity *Π* did not differ significantly between outlier and non-outlier gene regions (Table S8), indicating that there is no global reduction in genetic diversity in outlier gene regions. To assess how recombination rate correlates with candidate outlier gene regions, we used recombination rate estimates provided by Morgan et al. 2017. We found that recombination rates did not differ significantly between outlier and non-outlier gene regions (Table S8). Of 2118, 196 outlier genes occurred in a recombination coldspot (Figure 3), which did not correspond to an over-representation of outlier genes in coldspots of recombination (10,000 permutations, Z-score = −0.379, p = 0.663, Figure S2).

**Figure 3:**
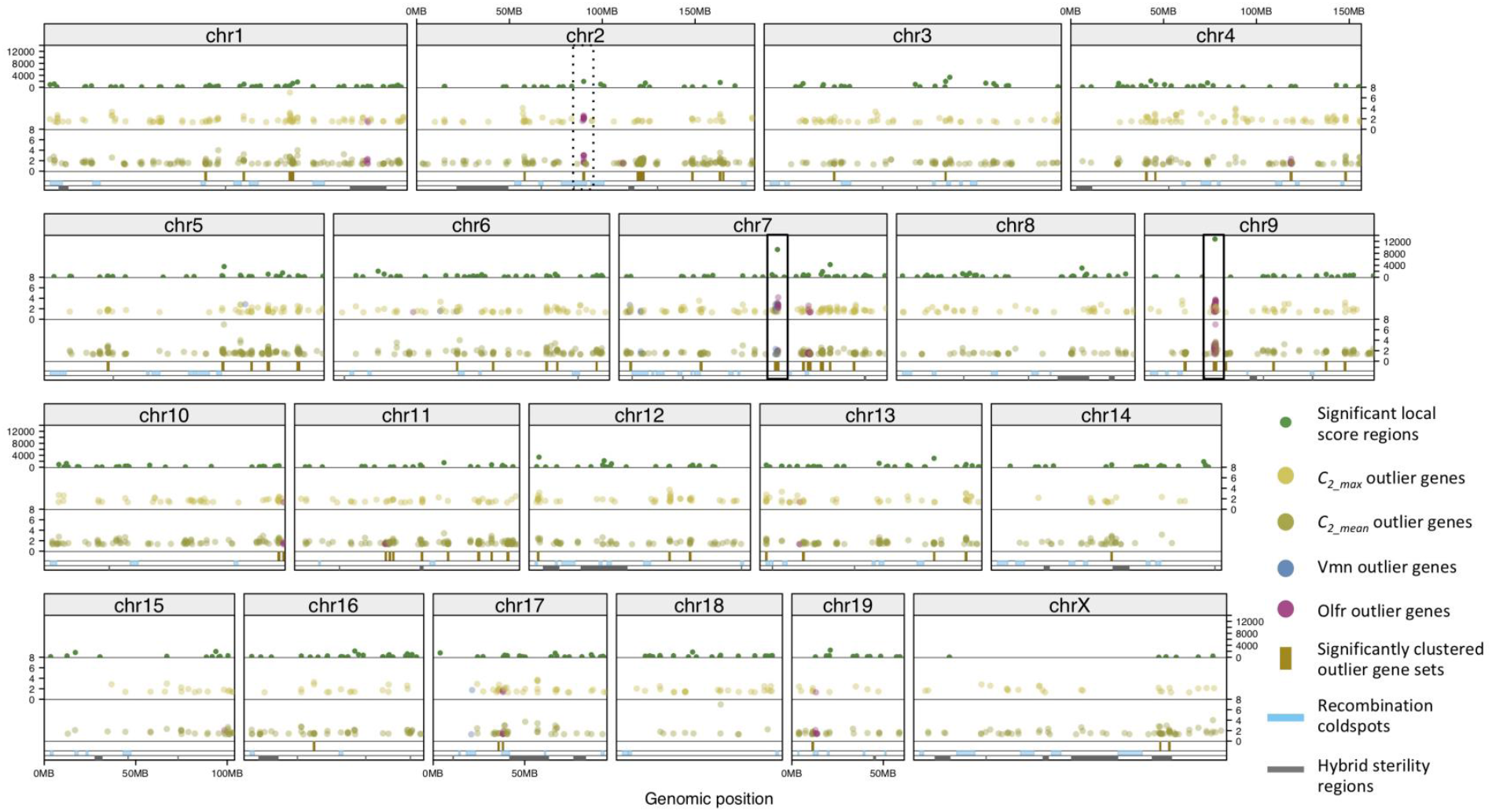
Genome-wide distribution of candidate regions showing significant genetic differentiation between Choosy and Non-Choosy populations. Track 1 (top): chromosomes; Track 2: local score values for significant genomic segments based on information on outlier SNPs; Tracks 3 & 4: *C_2_max_* and *C_2_mean_* values of outlier genes (922 and 1622 *C_2_max_* and *C_2_mean_* outlier genes shown respectively, over 41,702 annotated genes); vomeronasal (*Vmn*) and olfactory (*Olfr*) outlier genes are shown in blue and purple respectively; Track 5: significant outlier gene clusters; Track 6: recombination cold spots observed in the Diversity Outbred hybrid mouse population by Morgan et al. 2017; Track 7 (bottom): candidate hybrid sterility genes identified by Turner & Harr 2014. The three most significant outlier gene clusters are highlighted, on chromosomes 9, 7 and to some extent 2.

*SFS-based signatures of selective sweeps in genetic differentiation outliers :* to test whether differentiation-based outlier regions also show a signature of selective sweeps estimated from SFS-based methods, we run the program POOL-HMM (Boitard et al. 2013) on the two Choosy populations, the expected targets of reinforcing selection. We found 936 annotated genes within which at least one significant selective sweep window was detected in the Choosy1-Borum population, among which 28 *C_2_* outlier genes (including *Olfr291* on chromosome 7, and *Olfr1328*, *Olfr1339*, *Olfr1340* and *Olfr1342* on chromosome 4) (Table S9a). In the Choosy2-Låsby population, we found 1247 genes with at least one significant selective sweep window, among which 31 *C_2_* outlier genes (including *Olfr32*, *Olfr1258* and *Olfr1270* on chr2, and *Olfr62*, *Olfr1339*, *Olfr1340* and *Olfr1342* on chromosome 4) (Table S9b). However, only 3 *C_2_* outlier genes were found with significant selective sweep windows in both Choosy populations (expected under a reinforcement scenario). Those three genes are *Olfr1339, Olfr1340* and *Olfr1342*.

*No significant overlap between outlier genes and candidate hybrid sterility regions but some punctual associations exist:* we also tested whether outlier genes significantly co-localise with the 54 genomic regions identified as candidates for hybrid sterility (postmating isolation candidates) between *M. m. musculus* and *M. m. domesticus* (Turner and Harr 2014; Mukaj et al. 2020). We found that 143 outlier genes identified in our study overlap with candidate sterility regions (see Table S5 and Figure 3), which is not more than expected by chance (10,000 permutations, Z-score = −2.686, p = 0.997; Figure S3a). However, we found that out of the nine highly supported candidate sterility regions (i.e. showing concordant association with the two sterility phenotypes: relative testis weight and testis expression, see (Turner and Harr 2014)), 8 contained at least one outlier gene. The mean distance between outlier genes and sterility genomic regions was 23.70 Mb, which is more distant than expected by chance (10,000 permutations, Z-score = 3.929, *p* = 0.0001, Figure S3b). We note that PRDM9 gene, located on chromosome 17 and known as a major hybrid sterility gene in the house mouse (Mukaj et al. 2020), is distant by ~ 5 Mb (less than expected by chance, Figure S3b) from the closest outlier gene (*Vmn1r228*, see Table S5).

#### Clustered distribution of outlier loci in the genome

*Signals of outlier SNPs accumulate in two main genomic regions on chromosomes 9 and 7 :* data from 10-kb, 50-kb and 100-kb non-overlapping windows were examined, for each chromosome separately, for autocorrelation in genetic differentiation associated with the choosy/non-choosy trait, as measured by the BayPass *C_2_* contrast statistic. We found some significant autocorrelation and partial autocorrelation between neighbouring windows (Figure S4), with the autocorrelation parameter rho of a first-order autoregressive process estimated between 0.70 and 0.77 (e.g. 50-kb-windows, chr9: rho = 0.74, *p*-value = 0.039). These results indicated that there is a long-range dependence in genetic differentiation measures (BayPass *C_2_* contrast statistic) present over distances up to 100 kb. This dependence among *C_2_* measurements has a conservative effect on the identification of outlier SNPs: using only FDR and not an adjusted *α* based on the autocorrelation estimates, we corrected for an overinflated number of apparently independent tests, making the *p*-value cut-off conservative. Given this autocorrelation affecting SNP-based *C_2_* measurements, we applied the local score approach (Fariello et al. 2017) to identify outlier genomic segments cumulating signals from single SNP markers and characterize their distribution along the genome. This analysis identified 557 significant local score genomic regions, distributed on all chromosomes (Figure S5, Table S10 and Figure 3). Their mean size (+/- SE) is 99.139 kb (+/- SE: 6.528 kb). Two regions stood out as the most significant ones: one 1-Mb long region on chromosome 9 (peak value of the local score = 12769.03, coordinates: 9:37721717-38752579), containing 52 outlier genes among which 40 outlier Olfactory Receptor genes (Table S5); one 0.8 Mb long region on chromosome 7 (peak value of the local score = 9260.49, coordinates: 7:85585560-86388933), containing 17 outlier genes, among which 10 Vomeronasal Receptor and 5 Olfactory Receptor outlier genes (Table S5).

*Two ~ 2 Mb long Olfactory and Vomeronasal receptor gene clusters on chromosomes 9 and 7 represent the main candidate genomic regions underlying behavioural divergence*: using the expected number of genes per 500 kb and assuming a random genome-wide distribution, a permutation test showed that outlier genes were not homogeneously distributed along the genome (p = 2e-05, 50,000 permutations) (Figure S6). We, therefore, analysed this pattern of genomic clustering in more detail, by investigating whether some outlier gene sets were more clustered in the genome than expected by chance. We identified 63 outlier gene sets found significantly clustered in the genome (Table S11 and Figure 3). The most significantly clustered outlier gene sets were found on chromosome 1 (cluster #3, p = 9.37E-14), chromosome 5 (cluster #20, p = 7.31E-05), chromosome 9 (cluster #39, p = 7.31E-05; cluster #35 (Olfr cluster), p = 5.24E-04), chromosome 11 (cluster #49, p = 5.24E-04), chromosome 12 (cluster #52, p = 5.24E-04) and chromosome 7 (cluster #28 (Vmn+Olfr cluster), p = 2.97E-03). The mean size of those significant outlier gene clusters was 919.88 kb +/- SE: 66.10 kb. The largest gene clusters were found on chromosome 2 (cluster #6: 3.7 Mb), chromosome 1 (cluster #3: 2.45 Mb), chromosome 7 (cluster #28 (Vmn+Olfr cluster): 2.3 Mb; cluster #30 (Olfr cluster): 1.95 Mb) and chromosome 9 (cluster #35 (Olfr cluster): 1.8 Mb). These cluster sizes extend far beyond the estimated mean autocorrelation effect on each chromosome, present over distances up to 100 kb. Figure 3 recapitulates the distribution of significant local score regions, outlier genes including Olfr and Vmn ones, and significant outlier gene clusters along the genome.

By counting the total number of outlier and ‘top’ outlier genes and applying weights to this content (see Table S11 for details), we could identify the gene clusters showing the strongest degree of genetic differentiation between Choosy and Non-Choosy populations. Three gene clusters stood out as the most differentiated ones: gene cluster #35 on chromosome 9, gene cluster #28 on chromosome 7, and to some extent gene cluster #5 on chromosome 2 (Table S11). Strikingly, these 3 gene clusters correspond to the three main olfactory/vomeronasal receptor outlier gene clusters: cluster #35 contains 43 *Olfr* outlier genes; cluster #28 contains 14 *Vmn* and 6 *Olfr* outlier genes; cluster #5 contains 11 *Olfr* outlier genes. These three outlier gene clusters stand out from genome-wide patterns of differentiation (Figure 3, Figure S7) and the two most important ones on chromosomes 9 and 7 are detailed in Figure 4. Cluster #35 on chromosome 9 contains the five *Olfr* outlier genes identified among the 10 most differentiated ‘top’ outlier genes (Table 1 and Table S5).

**Figure 4:**
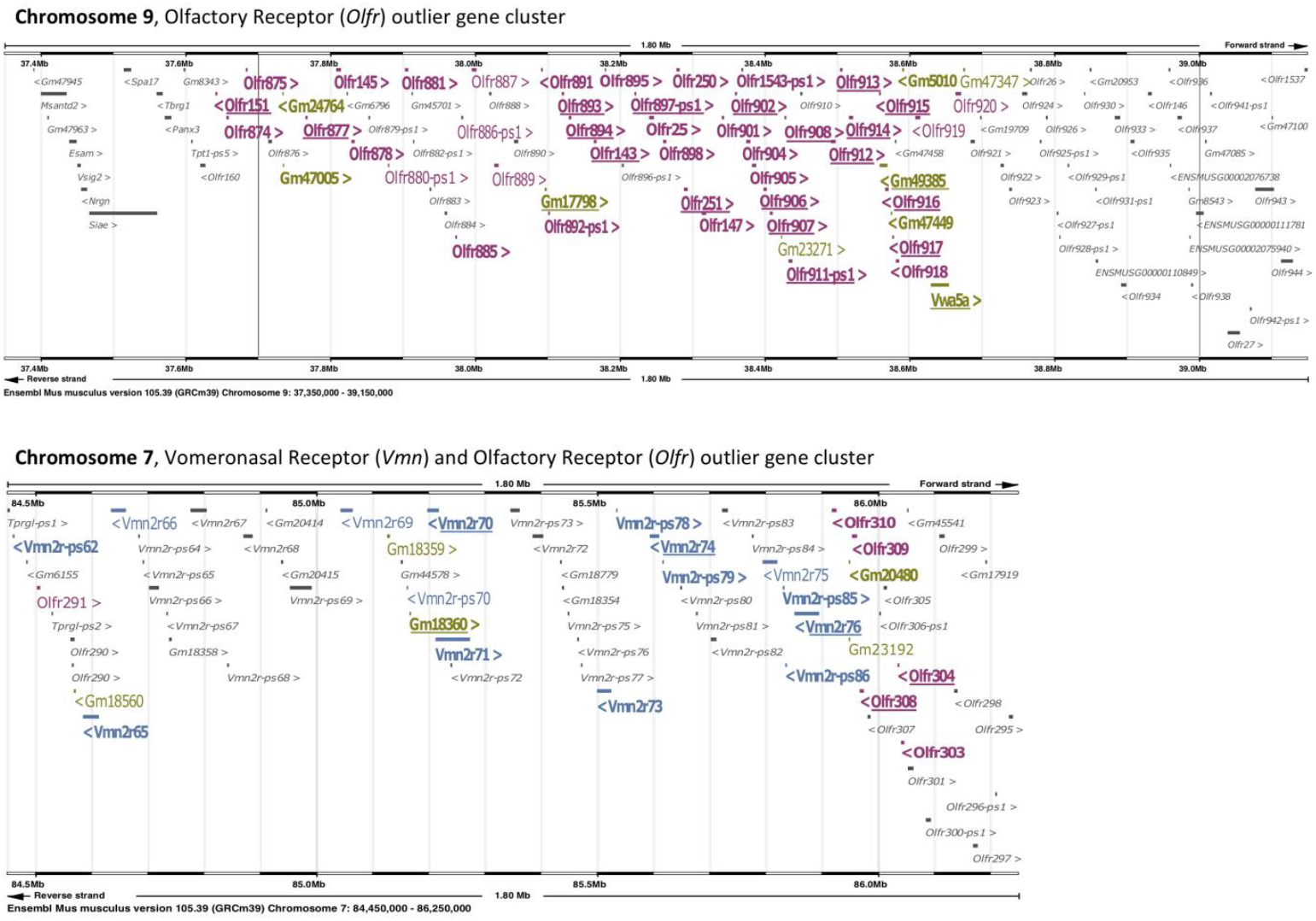
The two main candidate gene clusters composed of Olfactory and Vomeronasal Receptor genes on chromosomes 9 and 7. *Ensembl* protein-coding names and locations are shown within these two 1.8 Mb long genomic regions. The “>“ or “<“ symbols indicate the orientation of the gene on the chromosome. Non-outlier genes are in grey and outlier genes are in colours (purple: Olfactory Receptor genes; blue: Vomeronasal Receptor genes; green: any other outlier genes). For outlier genes in colours, different levels of significance are indicated as follows: regular text: *C_2_max_* or *C_2_mean_* outlier genes at *p* < 0.05; bold text: *C_2_max_* or *C_2_mean_* outliers at *p* < 0.01; bold and underlined: top significant outlier genes, based on the combined information from *C_2_max_* and *C_2_mean_* statistics.

## Discussion

In this study, we addressed the genetic basis of assortative mate preferences in the house mouse, expected to be evolving under the action of reinforcing selection, using a population genomics approach coupled with functional predictions. We identified some regions of the genome showing significant and consistent divergence between populations with contrasted behaviours (Choosy populations from the hybrid zone and Non-Choosy populations from allopatric areas). A series of Olfactory and Vomeronasal Receptor (OR and VR) genes, among the most differentiated regions and in line with functional predictions, stand out as the prime candidates potentially underlying this olfactory-based behavioural divergence. Those outlier genes form significantly clustered sets of genes in the genome, with two main candidate gene clusters on chromosomes 9 and 7, extending up to 1.8 Mb and containing several highly significant Olfactory and Vomeronasal receptor outlier genes.

### Population genomics and the genetics of complex trait divergence

Population differentiation scans, between two or more populations that differ in a behaviour trait of interest, have rarely been applied to address the genetic basis of polygenic behaviours (but see (Toews et al. 2019)). However, they can be powerful for mapping genetic variation at a high resolution, can be applied to natural populations, and can additionally provide evidence of selection (Niepoth and Bendesky 2020). Potential drawbacks to these approaches include their agnosticism to phenotypic traits and the challenges to interpret signals of genetic differentiation (e.g. populations often differ in more than one trait, other factors than divergent selection can drive genetic differentiation - Cruickshank & Hahn 2014; Ravinet *et al*. 2017; Wolf & Ellegren 2017). In this study, we took advantage of this powerful population genomics approach but combined it with specific experimental and data analysis strategies as well as clear functional predictions to overcome its potential limitations and to refine interpretations of the results in terms of genetic variation associated with the observed behavioural divergence.

Our strategy to investigate population differentiation, with whole-genome resequencing data, at different genomic scales (SNPs, genes, different gene-based statistics) enabled us to identify highly supported candidate regions found consistently divergent regardless of the scales and the statistics used, while also characterising more comprehensively the catalogue of genomic regions and architectures potentially underlying this behavioural shift. Genome-wide SNP-based differentiation analyses, despite their limitations (Huang et al. 2011), informed about the respective role of genic versus non-genic regions (mostly regulatory regions) in population differentiation, while confirming functional enrichment results obtained with gene-based analyses. However, our study confirmed that genome-wide gene-based differentiation analyses are efficient for gaining power and precision as well as for facilitating the link with the phenotype of interest (Neale and Sham 2004; Huang et al. 2011; Chung et al. 2019). The identification of enriched functional categories and the most promising individual genes was eased. Moreover, by capturing different potential underlying genetic architectures of divergence (single versus multiple small-effect causal variants within a gene) (Wang et al. 2017), the two different gene-based statistics (*C*_*2*_max_ and *C*_*2*_mean_) provided two lists of outlier genes partially overlapping but also complementary. For example, most Olfactory Receptor candidates are *C*_*2*_mean_ outlier genes, while most Vomeronasal Receptor ones are *C*_*2*_max_ outlier genes.

The use of the *C_2_* contrast statistic of BayPass enabled us to assess genetic differentiation based on contrasts of allele frequencies corrected for population structure (Olazcuaga et al. 2021). The combination of biological replicates per behaviour group and multiple population comparisons (Focal and Control tests) allowed us to identify the genomic regions consistently differentiated between Choosy and Non-Choosy populations, beyond population-specific differentiation (Figure 1). These multi-way comparisons across populations had been advocated as a promising way forward to separate processes acting similarly on all populations from processes that are specific to a single population or population comparison (Wolf and Ellegren 2017) but are rarely implemented (but see Roesti *et al*. 2014; Vijay *et al*. 2016). In our study, this procedure narrowed down the list of genomic regions of interest, the control test removing respectively 26% and 7.65% of the outlier SNPs and genes, with the final sets of outlier SNPs and genes eventually representing 0.017% and ~5 % of the total number of SNPs and annotated genes in the dataset. Yet, this corresponds to a relatively large number of outlier SNPs and genes (4818 and 2118 respectively) that show the expected patterns of genetic differentiation associated with behavioural divergence and reinforcement.

Importantly, we do not expect all of these differentiated genomic regions between Choosy and Non-Choosy populations to be involved in the observed behavioural divergence nor to be subject to reinforcing selection. Because these sets of populations can differ in more than one trait and selective regime, some differentiated regions may not affect the behaviour of interest. Moreover, neutral clines across the *M. m. domesticus/M. m. musculus* hybrid zone can produce genetic structure at specific loci between Choosy populations and Non-Choosy populations and hitchhiking effects can induce genetic differentiation at variants in the vicinity of causal variants. By selecting populations that are known to show distinct mating preferences despite being closely related and inhabiting a similar environment (in Jutland Denmark, separated by no more than 50 km), we minimized the risk of confounding selective effects, but the list of outlier loci extends very likely beyond the variants directly associated with the behavioural divergence. One key step of this study was therefore to confront genome scan results with functional predictions, here available from a series of previous studies that had characterised the behaviour of interest in detail (Smadja and Ganem 2002; Smadja et al. 2004; Smadja and Ganem 2005; Ganem et al. 2008). Matching very clear functional analyses’ results (functional enrichment, nature of outlier genes), indicating a prominent role of some olfactory and vomeronasal receptor genes, with information on the involvement of olfactory-based processes in the expression of assortative mate preferences, provides very strong support for considering these highly significantly differentiated olfactory and vomeronasal receptor genes as the prime candidates potentially associated with behavioural divergence and reinforcing selection (Table 1). Since our previous studies had shown that individuals from Non-Choosy populations can discriminate between mating signals of the two subspecies, but do not establish a directional mate choice (Smadja and Ganem 2005; Smadja and Ganem 2008), the genomic regions identified in our study likely underlie the evolution of choosiness rather than discrimination. Our study illustrates the importance of integrating detailed phenotypic characterisation with genomic data to gain power and accuracy in population genomic approaches that tackle the genetics of complex trait divergence.

### Genetic architecture of candidate regions and selective regimes

Autocorrelation analyses indicated a long-range dependence in genetic differentiation measures present over distances up to 100 kb, consistent with previous estimates of background linkage disequilibrium in wild house mouse genomes (Laurie et al. 2007; Staubach et al. 2012) and that could inflate the number of outlier loci (Hahn 2006). However, local score analyses (Fariello et al. 2017), which take into account this autocorrelation and background linkage disequilibrium, consistently pointed to numerous outlier local score genomic segments spread across several chromosomes (Figure 3). Whatever the criteria we applied (significance, consistency across analyses and statistics) to stringently narrow down the list of candidate genes and genomic regions, evidence supports a highly polygenic basis underlying the mate preference shift. From the initial list of 2118 outlier genes containing 127 Olfactory receptor (*Olfr*) and 27 Vomeronasal receptor (*Vmn*) genes, the most restricted set of ‘top’ outlier genes still include 95 genes spread across several chromosomes and among which 23 are *Olfr* and 4 are *Vmn* genes. A highly polygenic basis for assortative mate preferences fits predictions from a previous phenotypic cline analysis in the house mouse suggesting the involvement of a polygenic autosomal system with dominance and epistasis (Ganem et al. 2008). This result is also in line with the few studies addressing the genetics of variation in olfactory-mediated behaviours in insects (Swarup et al. 2013; Arya et al. 2015) and mammals (Santos et al. 2018).

Given the highly polygenic nature of genetic architecture underlying the behavioural shift and the apparent strong linkage disequilibrium among genes involved in this phenotypic divergence, candidate loci are likely under polygenic selection, expected for such complex traits (Latta 1998; Sella and Barton 2019). Interestingly, previous studies have shown that polygenic adaptation can play a major role in the evolution of species-specific innate behaviours through the modifications of pre-existing genetic networks (Anholt 2020) and in the rapid evolution of chemosensory systems (Librado and Rozas 2016). The fact that we did not find signatures of selective sweeps using SFS-based methods in most candidate outlier regions may be another indication of polygenic adaptation, known to be more detectable with pairwise tests for allelic differentiation (Stölting et al. 2015; Schneider et al. 2021). This result, together with the absence of a significant reduction of genetic diversity in outlier regions, may also suggest that those candidate regions are more likely subject to soft sweeps than hard sweeps (Pritchard et al. 2010; Wang et al. 2022). The two types of outlier genes (*C*_*2*_max_ and *C*_*2*_mean_) may here not reflect different selection regimes but just different variant architectures within outlier genes. Since outlier gene regions are not associated with regions of reduced genetic diversity or reduced recombination rate, a general effect of background selection inflating the levels of genetic differentiation in our data seems unlikely. However, future analyses on individual data and a more extensive sampling of Choosy populations would be an interesting perspective to get fine-scale estimates of recombination rate along these genomes and more directly test for signatures of polygenic selection/soft sweeps (Kern and Schrider 2018).

Another striking result is that outlier loci showed consistent signals clustered distributions in the genome. While the signals of outlier SNPs via local score analyses accumulated in two main genomic regions on chromosomes 9 and 7, clustering analyses of outlier genes consistently identified numerous sets of genes significantly clustered in the genome, amongst which the two most significant clusters pinpointing the same candidate regions on chromosomes 9 and 7. These two clusters of outlier genes contain several candidate olfactory and/or vomeronasal receptor genes and extend up to 1.8 Mb (Figure 4). These patterns suggest strong linkage disequilibrium in these large candidate regions, potentially indicative of the strength of selection acting on its targets or of complex correlations among loci belonging to the same gene regulatory network (Fagny and Austerlitz 2021). Within these clusters, it is difficult to say which genes and nucleotides are the direct targets of selection and pinpointing individual genes’ contributions to the observed behavioural shift in these genomic clusters will require additional experiments. Nevertheless, some outlier genes are more significantly differentiated than others in those clusters and can be considered prime candidates (*Vmn2r70, Vmn2r74, Vmn2r76, Olfr304* and *Olfr308* in the cluster on chromosome 7; *Olfr143, Olfr906, Olfr907, Olfr908* and *Olfr914* in the cluster on chromosome 9).

The source of this genetic variation specific to the *M. m. musculus* Choosy populations is still unknown. Considering our results, it may originate from standing variation present in allopatric populations or differential introgression of *Mus musculus domesticus* alleles across the hybrid zone. The large genomic clusters identified as outlier regions could fit the second scenario, which echoes the recent evidence of adaptive introgression of OR and VR gene clusters across species boundaries in the *Mus* genus (Ullrich et al. 2017; Baird et al. 2022; Banker et al. 2022). This scenario would be compatible with the role of these genomic regions in reproductive isolation with the nearby subspecies *M. m. domesticus*: the selective pressure of reinforcement could have favoured the adaptive introgression of alleles conferring choosiness on either side of the hybrid zone (a case of one-allele mechanism), the direction of the choice being possibly expressed via a matching rule, i.e. towards a mating partner matching the chooser’s genomic background (or mating signal type). These mechanisms are theoretically predicted to favour the evolution and maintenance of the barrier trait in the context of gene flow (Kopp et al. 2018). Additional genomic data for *M. m. domesticus* populations from the other side of the hybrid zone in Denmark would be needed to first test if variation comes from introgression.

Overall, these results on the genetic architecture and selective regimes associated with the divergence in mate preferences correspond to theoretical predictions for loci involved in adaptation and reinforcement. Concentrated architectures of adaptive loci are expected in the presence of gene flow (Yeaman and Whitlock 2011; Yeaman 2013), and fit with the few other empirical studies testing for this pattern of clustering in natural systems (e.g. (Fustier et al. 2017; Roda et al. 2017). Moreover, the selective regime of reinforcement corresponds to indirect selection on prezygotic isolating traits in hybrid zone populations, usually from standing genetic variation present in allopatry (Felsenstein 1981; Kirkpatrick and Servedio 1999) via linkage disequilibrium between loci under direct selection (hybrid sterility genes) and assortative mating loci. We were expecting a co-localisation between candidate hybrid sterility genomic regions (Turner and Harr 2014) and outlier regions that could facilitate the maintenance of linkage disequilibrium among those co-adapted traits (mate preference and hybrid sterility). With the exception of a few co-localising regions, we did not find any significant physical overlap between these two sets of genes. However, this test may miss some unidentified hybrid sterility genes and is quite stringent (testing overlap and not physical linkage). For example, the outlier gene cluster on chromosome 9 lies in physical proximity with a candidate hybrid sterility region (Figure 3). Moreover, linkage disequilibrium can still evolve and be maintained between unlinked or loosely linked and is not necessary under the scenario mentioned above of a case of one-allele mechanism (Servedio 2009; Butlin and Smadja 2018; Hench et al. 2019). Our study provides an important addition to the rare studies that have investigated the genetics of reinforcement (Ortiz-Barrientos et al. 2004; Hopkins and Rausher 2011; Ospina et al. 2021), by testing predictions on expected genomic patterns of differentiation to characterize the genetic basis of a complex premating isolating trait and the genomic signatures of reinforcing selection acting on it.

### The role of Olfactory and Vomeronasal Receptor genes in the evolution of mate choice divergence

Chemoreceptor genes are one of the most extensive and diverse multigene families in most vertebrate and non-vertebrate genomes (Sánchez-Gracia et al. 2011; Ibarra-Soria, M. Levitin, et al. 2014; Benton 2015; Bear et al. 2016). Yet, their precise role in key evolutionary processes such as adaptation and speciation remains largely unknown, although chemosensory processes have been identified as key players at the phenotypic level (Smadja and Butlin 2009). Comparative analyses have provided great insights into the role of OR or VR repertoire evolution in diversification and ecological adaptations (e.g. dietary specialisation (Hayden et al. 2014; Yohe et al. 2021) or adaptation to different life-styles (Shi and Zhang 2007; Hayden et al. 2010; Khan et al. 2015; Jiao et al. 2019). At microevolutionary scales, studies are scarce due to the difficulty of analysing these gene families and linking their genetic variation to relevant adaptive phenotypes and selective regimes (but see (Smadja et al. 2012; Li et al. 2015; Poelstra et al. 2018; Auer et al. 2020). Our study provides one of the clearest and most detailed examples of the involvement of olfactory and vomeronasal receptors in the evolution of an adaptive behavioural trait, assortative mate choice, and how natural selection drives their genetic divergence.

Our results indicated the highly significant enrichment of this category of receptor genes among outlier genomic regions (Figure 2B), in agreement with our functional prediction.

General variant effect predictor analyses pointed to the possible dual role of regulatory and proteincoding changes in the evolution of behavioural divergence, with the presence of missense and stop gained variants as well as variants in already known regulatory regions or putative regulatory regions in downstream / upstream gene regions (Figure 2A). For candidate *Olfr* and *Vmn* genes, many SNP variants fall within upstream or downstream gene regions, UTR regions or intronic regions potentially affecting gene regulation, but we identified a few protein-coding changes affecting *Olfr250, Olfr906, Olfr908, Olfr915, Olfr916* in the candidate cluster on chromosome 9, and *Olfr304* and *Vmn2r76* in the cluster on chromosome 7 (Table 1). Consistent with a putative role of regulatory changes are the results of two of our previous studies investigating the expression and copy-number divergence between the same Choosy and Non-Choosy populations. We found significant differential expression at some Vomeronasal Receptor genes expressed in the vomeronasal organ (Loire et al. 2017), four out of five (*Vmn2r69, Vmn2r70, Vmn2r73, Vmn2r75*) being among the *Vmn* outlier genes identified in the present study and all belonging to the promising candidate gene cluster on chromosome 7. We also identified deletions specific to the Choosy populations in upstream or intronic regions of some Olfactory Receptor genes (*Olfr907* in the candidate cluster on chromosome 9, *Olfr301*, in the candidate cluster on chromosome 7, *Olfr564* and *Olfr461* on chromosomes 7 and 6), that could affect their expression or the expression of nearby *Olfr* genes (North et al. 2020). The combination of nucleotide, structural and expression variation primarily at Olfactory and Vomeronasal Receptor genes could therefore shape the evolution of this complex behavioural trait under selection (Niepoth and Bendesky 2020; Fagny and Austerlitz 2021).

The Olfactory and Vomeronasal Receptor prime candidates are members of some specific phylogenetic clades of closely-related receptors within each gene superfamilies. This result echoes previous insights obtained by Isogai *et al*. on the functional characterisation of Vomeronasal Receptors in the house mouse (Isogai et al. 2011). They showed that distinct mouse VR receptor subfamilies have evolved towards the specific recognition of certain animal groups (heterospecific signals from predators or sympatric closely related species/subspecies, male or female conspecific signals) or chemical structures (volatile steroids, non-volatile proteins). Interestingly, some members of V2R clades 3 and 6 (our main VR candidate clusters) were shown to be involved in the recognition of closely-related and sympatric species or conspecifics of the opposite sex, leading the authors to suggest that “through the activation of specialized receptors, *M. musculus* may readily discriminate scents emitted by closely related but reproductively incompatible species, a property that could be linked to the reproductive isolation of these species”. Considering the results of our study, and if we extrapolate Isogai’s results to ORs, we can hypothesize a specialization of some VR and OR gene clusters towards the function of assortative mating in the hybrid zone.

In the future, several interesting follow-up from our work will help to better understand the role of these candidate receptors in the evolution of assortative mating. Comparative transcriptomics on the main olfactory epithelium will complement the study carried out on VNO (Loire et al. 2017) to assess whether changes in expression affect *Olfr* genes between Choosy and Non-Choosy populations. Haplotagging (Meier et al. 2021) and long-range sequencing will help to further test for polygenic selection and structural changes in these candidate genomic regions. Finally, functional characterisation of the candidate OR and VR gene clusters could be tackled by leveraging the molecular tools developed by Isogai et al. (i.e. activity-dependent immunolabelling of sensory neurons coupled with *in situ* hybridisation) and applying them to wild mice from hybrid zone. With this approach, we could test whether those receptors are activated by subspecific cues (i.e. urinary stimuli), as expected if involved in the expression of assortative mate preferences.

## Methods

### Samples

We trapped *M. m. musculus* wild adult mice in Jutland, Denmark in several sites (indoor farms and other human dwellings) and maintained them in the laboratory under controlled conditions before being behaviourally tested and euthanized for dissection. Sampling sites represented two distinct geographical areas in Jutland, which are characterised by populations with distinct mate preference behaviours but sufficiently geographically close to avoid any geographical effect of distant allopatry: (1) the border of the hybrid zone on the *M. m. musculus* side (50 km North to the genetic centre of the hybrid zone (defined in Raufaste et al. 2005)) where strong assortative mating was previously documented (Smadja et al. 2004; Smadja and Ganem 2005; Ganem et al. 2008) and confirmed on these newly sampled populations (Latour et al. 2014; Smadja et al. 2015) (‘Choosy’ populations), (2) another area in Jutland but further north from the hybrid zone, where a previous study did not find any significant directional mate preference in these sampled populations (‘Non-Choosy’ populations) (Latour et al. 2014; Smadja et al. 2015).

Two populations per geographical area were included as biological replicates, each composed of several trapping sites (four populations in total, hereafter called « Borum Choosy1 », « Låsby Choosy2 », « Hobro Non-Choosy1 » and « Randers Non-Choosy2) (see Figure 1a and Table S1) for a detailed sample description). For all individuals from each of the four populations *n*_Borum_ = 33, *n*_Låsby_ = 27, *n*_Hobro_ = 29, *n*_Randers_ = 29), spleen was extracted rapidly after death by cervical dislocation, immersed in ethanol and stored at −80°C. We extracted genomic DNA from individual mouse spleens, using the Macherey-Nagel kit standard protocol.

### Data generation and variant processing

#### Whole-genome re-sequencing

To assess genomic divergence in two independent comparisons of Choosy and Non-Choosy populations, at the level of the whole genome (genome size: 2.7 Gb), we carried out a pool-sequencing approach. Pooled samples representing each population were obtained by mixing individual DNA extracts from each population in equimolar proportions. Library preparation and sequencing were carried out at the MGX-Montpellier GenomiX platform (Institut de Génomique Fonctionnelle - Institut de Génétique Humaine, Montpellier, France). Sequencing libraries (average insert size: 478 bp (Borum Choosy1), 497 bp (Låsby Choosy2), 478 bp (Hobro Non-Choosy1) and 445 bp (Randers Non-Choosy2)) were prepared using two methods to increase sequence diversity: Illumina TruSeq DNA Sample Prep Kit v2 (three-quarters of samples), and Illumina Nextera DNA sample preparation kit. Overall, 3/4 of all sequence data was obtained using the TruSeq protocol. All libraries were sequenced on an Illumina HiSeq2000 machine using a paired-end 100-bp read-length protocol, targeting an average raw coverage per pool of 30X, chosen as a cost-effective optimum given the large house mouse genome and pool size/pool coverage balance recommendation for pool-seq sequencing (Futschik and Schlotterer 2010; Gautier et al. 2013; Schlötterer et al. 2014). Sequencing reads were trimmed and cleaned of adaptor sequences using Trimmomatic (Bolger et al. 2014).

#### Mapping, SNP identification and filtering

We applied stringent and optimized mapping and SNP filtering conditions to obtain accurate allele frequency estimations from pool-seq data (Gautier et al. 2013; Schlötterer et al. 2014). The program Bowtie2 v2.2.3 (Langmead and Salzberg 2012) was used to map the reads against the house mouse reference genome (http://www.ensembl.org/, version GRCm38.p2), with the –very-sensitive-local option to get a stringent alignment and a fragment size limit of 600 (-X parameter) to increase the number of concordant mapping count and reduce the discordant mapping count. The Samtools suite v1.3.1 (Li et al. 2009) was used to sort, index and filter for read duplicates the resulting BAM files. An overview of sequencing and mapping statistics can be found in Table S2. BAM files were combined into mpileup files and *sync* files (using samtools mpileup and mpileup2sync.pl script from Popoolation2 package (Kofler et al. 2011)) and first filtered based on base quality (min-qual 20). We filtered out *sync* files for indels using Popoolation2, and developed a custom Python script to further filter bi-allelic SNP variants based on minimum coverage (= 10X), maximum coverage (= 100X, to avoid repeated/duplicated regions of the genome), and minimum count of minor allele (= 3). Minimum and maximum coverage filters were applied per population, and a minimum allele count filter was applied across all populations. From this set of filtered SNPs, we estimated mean population genetic diversity *Π* (NPSTAT package v1, Ferretti et al. 2013) and pairwise-population *F_ST_* (Pool-Seq estimator implemented in the *poolfstat* R package, Hivert et al. 2018; Gautier et al. 2022) to describe the populations under study (Figure 1a).

### Identification of SNPs and genes associated with behavioural divergence and under divergent selection

#### Population contrasts and BayPass procedure

We used the program BayPass v2.2 (Gautier 2015) to identify Single-Nucleotide-Polymorphisms (SNPs) significantly and repeatedly divergent between Choosy and Non-Choosy populations, i.e. associated with the observed behavioural divergence between these populations. BAYPASS implements a method accounting for the neutral correlation of allele frequencies across populations to detect loci under selection or associated with covariables. We used BAYPASS to estimate the scaled covariance matrices (Ω) of the population allele frequencies for autosomal and X-linked SNPs (Figure 1b). This matrix captures information about the shared demographic history of the sampled populations which is expected to differ between autosomes and the X chromosome (Clemente et al. 2018). Using our repeated experimental design (two biological replicates per behavioural group), we relied on the recently implemented *C_2_* statistic of BAYPASS (Olazcuaga et al. 2021) to contrast SNP allele frequencies corrected for the global effect of the shared population history captured by the matrix Ω on their covariance (i.e. standardized allele frequencies) between the two groups of populations specified by the behavioural trait. This approach allows correcting for population structure to identify SNP associated with binary co-variables characterizing the populations under study which here consisted of two distinct comparisons (Figure 1c). First, the Focal Test consisted in estimating the contrast statistic between the two groups of populations with contrasted mating behaviours: Choosy1+Choosy2 versus Non-Choosy1+Non-Choosy2 populations. This test aimed at identifying SNPs showing significant association with the behavioural contrast between Choosy and Non-Choosy populations. To further remove outlier loci from the Focal Test that would not show a consistent divergence between both Choosy and both Non-Choosy populations, we carried out a Control Test. This control test estimates the contrast statistic between two other combinations of the same populations, this time not grouped by their behavioural profile (one Choosy and one Non-Choosy population per group) (Figure 1c). We used the inference of genetic relationships among populations based on the Ω matrix estimated by BayPass (see Figure 1b) to maximize the distance between the control groups (i.e. comparing Choosy1+Non-Choosy2 versus Choosy2+Non-Choosy1) to perform the most stringent control contrast. SNPs and genes found to be significantly divergent in the Focal Test but also found to be significantly divergent in the Control Test (potentially reflecting population-specific divergence in the Focal Test) were removed from the final list of candidate loci.

Custom scripts were used to format filtered *sync* files into BayPass input files (Smadja et al. 2022b). We run BayPass separately for the autosomes and the X chromosome and specified the pool haploid sample sizes to activate the Pool-Seq mode of BayPass. For the autosomes, the pool haploid sizes were simply based on sample sizes (*n*__aut_Borum-Choosy1_ = 66; *n*__aut_Låsby-Choosy2_ = 54; *n*__aut_Hobro-NonChoosy1_ = 58; *n*__aut_Randers-NonChoosy2_ = 58). For the X chromosome, we adapted the haploid pool size information to the effective sex ratio of each pool (Table S1; *n*__chrX_Borum-Choosy1_ = 48; *n*__chrX_Låsby-Choosy2_= 41; *n*__chrX_Hobro-NonChoosy1_ = 45; *n*__chrX_Randers-NonChoosy2_ = 43). We used the BayPass core model with default options for the MCMC algorithm to obtain estimates of the scaled covariance matrix (Ω) and the *C_2_* contrast statistic. Following the BayPass manual instructions, the autosomal dataset was divided into sub-datasets of 110,905 SNPs (by taking one SNP every 250 SNPs along each chromosome) and three independent runs (using option - seed) were performed to assess the reproducibility of the MCMC estimates. We found a fairly high correlation across the different independent runs for the different estimators (Pearson’s *r* > 0.95 for the *C_2_* estimator) and thus only presented results from the first run. Similarly, a near-perfect correlation of the posterior means of the estimated Ω matrix elements was observed across independent runs as well as within each run across SNP subsamples, with the corresponding FMD distances (Gautier 2015) being always <0.4. We thus only reported BayPass results that were obtained from a single randomly chosen sub-dataset analysed in the first run.

#### Identification of outlier SNPs

For both the Focal and Control Tests, we computed SNP-based *p*-values from the *C_2_* values, the *C_2_* statistic following a Chi-squared distribution with one degree of freedom under the null hypothesis of no association (Olazcuaga et al. 2021). We identified outlier SNPs in the Focal Test (Choosy versus Non-Choosy) after correcting for multiple testing (*FDR < 0.05*). Only outlier SNPs that were not significantly differentiated in the Control Test (*p*-values > 0.05 without correction for multiple testing) were retained for further analyses.

#### Identification of outlier genes

Gene annotations were obtained from Ensembl v90 (Mus_musculus.GRCm38.90.geneset). Using SNP-based *C_2_* estimates and the BEDTOOLS suite v2.16.1 (Quinlan and Hall 2010) (*bedtools map*), we computed for each annotated gene in the house mouse genome two gene-based differentiation statistics: *C_2___max_* and *C2_mean*, corresponding to the highest SNP-based *C_2_* value of a gene and the average *C_2_* value of a gene, respectively. These two gene-based statistics correspond to two groups of gene-based methods: *C_2_max_* has the advantage of detecting genes with a single main causal variant while reducing the total number of tests, while *C_2_mean_* appropriately captures the combined effect of multiple small-effect SNP variants within a gene (Wang et al. 2017). Since we had no *a priori* knowledge of the genetic architecture of our phenotype of interest (mate preference), nor on which one of the test statistics was optimal for our analysis, we used those two different statistics and reported as outlier genes those showing significant *C_2_max_* or *C_2_mean_* statistics. The significance of each gene-based differentiation statistic was assessed using custom permutation procedures (Smadja et al. 2022b). *C_2_max_* estimates are influenced by the nucleotide diversity of each gene: the larger the number of SNPs it contains, the more likely are extreme *C_2_max_* values. Thus, our permutation procedure consisted in sampling SNP-based *C_2_* values among gene sequences (including introns) in the genome equal to the number of SNPs in the focal gene. We then recorded the maximum *C_2_* value of the sample and generated a random distribution specific to the tested gene by repeating the resampling 10,000 times. *C_2_mean_* estimates are further influenced by gene length, a proxy for the degree of linkage disequilibrium among SNPs occurring in a given gene. To control for polymorphism and gene length in this permutation procedure, we thus sampled, for each gene, genomic fragments which satisfied two conditions: i) the same number of SNPs and ii) a length in base pairs not deviating from more than 10% of the observed one. For each fragment that satisfied these conditions (and thus mimicked the genomic properties of the gene), we calculated *C2_mean*. Repeating this process 10,000 for each gene allowed us to estimate an empirical *p*-value for the observed gene-based *C_2_mean_* statistic. Significant genes (for *C_2_max_* or *C_2_mean_* ) in each population comparison (Focal, Control) were therefore those with *p*-values < 0.05 or 0.01 (see below).

Only outlier genes that were significantly differentiated in the Focal comparison (*p*-values < 0.05) and not significantly differentiated in the Control comparison (*p*-values > 0.05) were retained. We defined as the final and complete list of outlier genes the union of *C_2_max_* and *C_2_mean_* outlier genes so that *C_2_max_* specific and *C_2_mean_* specific outlier genes are taken into account. All downstream analyses were performed on this set of outlier genes (with two stringency categories: *p*-values < 0.05 or *p*-values < 0.01). However, we also defined a restricted list of outlier genes (hereafter called ‘top outlier genes’), identified as highly differentiated based on the combined information from the two *C_2_max_* and *C_2_mean_* contrast statistics. To implement this classification, we first assigned each outlier gene to a significance category for each gene-based statistic (1: most significant; 5: least significant), assigned weights to these categories and calculated an overall differentiation score as the sum of the weights obtained from *C_2_max_* and *C_2_mean_* information. Genes were classified as ‘top outlier genes’ when showing a minimum overall differentiation score of 10.

### Functional and genomic characteristics of candidate genomic regions

#### Functional analyses of candidate genomic regions

The final set of outlier SNPs inferred from the combination of the Focal and Control BayPass Tests was analysed using the Ensembl Variant Effect Predictor (VEP) (McLaren et al. 2016) to annotate outlier SNPs in coding and non-coding genomic regions as well as to determine their impact on the protein sequence or gene regulation. We used the program David v6.8 (Huang et al. 2009) to carry out functional annotation analyses on both outlier SNP and gene sets. Enriched annotation terms were identified using a significance threshold of 0.01 (EASE score) after Benjamini & Hochberg’s correction. To reduce the functional redundancy among annotation terms, we also carried out Functional Annotation Clustering analyses provided by David, which report functional groups with similar annotation terms (significant enrichment score (i.e. the geometric mean of all the enrichment *p*-values (EASE scores) for each annotation term associated with the gene members in the group) > 1.3). Over-representation of some candidate gene families such as Olfactory Receptor (*Olfr*) and Vomerosanal Receptor (*Vmn*) genes among outlier gene sets was tested using hypergeometric tests (*phyper* R function).

#### Genomic features associated with outlier gene regions

We used the NPstat package v1 (Ferretti et al. 2013) to estimate nucleotide diversity Π (Tajima 1983), measured as the average pairwise nucleotide diversity for a given position across all reads and summing it over all positions, and Watterson’s theta θ (Watterson 1975) statistics in 5 kb and 50 kb non-overlapping windows across all 20 chromosomes for each of the four studied populations. Using the BEDTOOLS (bedtools flank and map) and custom scripts, we computed nucleotide diversity Π for all annotated gene regions (gene sequence +/- 5 kb) and tested for differences between outlier and non-outlier gene regions using Kruskall-Wallis tests. Pool-hmm v1.4.3 program (Boitard et al. 2013) was used to detect selective sweep windows across the genome in each of the two Choosy populations, using estimates of Watterson’s θ provided by NPstat.

We tested for non-random co-localization of candidate outlier genes identified in this study with (i) coldspots of recombination reported by Morgan et al. 2017 and (ii) candidate hybrid sterility genomic regions between *M. m. musculus* and *M. m. domesticus* identified by Turner and Harr 2014. Candidate hybrid sterility regions correspond to the 53 GWAS genomic regions significantly associated with relative weight testis variation (named RTW regions) and/or testis expression variation (named PC regions) (Turner and Harr 2014). We added to this list of candidate genes the gene *PRDM9* (chr17:15543079-15564354) known as a major hybrid sterility gene in the house mouse. We counted the number of outlier genes overlapping these two genomic region sets (coldspots of recombination and candidate sterility regions) and used permutations to assess whether outlier genes overlap significantly more than by chance with these genomic elements (outlier genes resampled 10,000 times among all annotated genes). To perform these permutation tests, we used the bioconductor *RegioneR* (Gel et al. 2015) package in R version 3.6.2 (R Core Team 2019). We also performed a Kruskall-Wallis test to investigate whether recombination rates differ significantly between outlier and non-outlier gene regions.

#### Genomic distribution of outlier SNPs and genes

*Outlier SNPs*: we first estimated the autocorrelation between linked loci in the level of genetic differentiation (BayPass SNP-based contrast statistics *C_2_*) using a first-order autoregressive model (Hahn 2006). For each chromosome, 10 kb, 50 kb and 100 kb non-overlapping windows were analysed for autocorrelation and partial autocorrelation in *C_2_* (using the *acf* and *pacf* R functions respectively). We then applied the local score approach (Fariello et al. 2017), which accounts for linkage disequilibrium using autocorrelation information, to identify outlier genomic segments cumulating signals from single SNP markers and characterize their distribution along the genome. We looked at the distribution of -log_10_(*p*-value) to determine the tuning parameter *ξ*, which must be comprised between mean(-log_10_(*p*-value)) (here equal to 0.425) and max(-log_10_(*p*-value)) (here equal to 10.365). We chose a value of *ξ*= 1 to run the excursions of the Lindley process so that we could expect cumulating a larger number of intermediate signals and the identification of wider outlier regions, putting more emphasis on recent soft or incomplete sweeps. Although this method does not provide absolute information about the number and the size of candidate genomic regions since they depend on the tuning parameter *ξ*, it is useful to identify regions with outstanding genetic differentiation between populations and the genomic range at which cumulating effects at multiple differentiated SNPs spread.

*Outlier genes*: we first assessed whether outlier genes were distributed heterogeneously in the genome (permutation procedure testing whether the observed number of outlier genes per 500 kb was larger or smaller than expected by chance). We then performed a cluster analysis (developed by (Haque et al. 2016)) to identify the number, location and size of outlier gene clusters in the genome. For the clustering approach, the window size used was 500 kb (basal autocorrelation being non-significant at distance > 100 kb) and this 500-kb window was shifted 50 kb at a time in an overlapped fashion. For each sliding window, the number of outlier genes was counted. The sliding window was used from the start of the chromosome until the end for each chromosome separately. Once the number of outlier genes had been calculated for each window a *z* -test was used to calculate the *z* -score for each window from which the *p*-value was calculated. Any window with a *p*-value < 0.05 was taken as statistically significant in the over-representation of outlier genes. Windows falling within 50,000 bases (consecutive windows) of each other were merged to create final clusters. We graphically represented the distribution of significant local score segments, outlier genes and outlier gene clusters along the genome using the Gtrellis v1.4.2 package (Gu et al. 2016).

## Acknowledgements

Preprint version 3 of this article has been peer-reviewed and recommended by Peer Community In Evolutionary Biology (https://doi.org/10.24072/pci.evolbiol.100157). We would like to thank very much Josette Catalan, Marco Perriat-Sanguinet, Yamin Latour for their help in collecting mice in Denmark, the Danish farmers for their hospitality, as well as Giveskud zoo personnel. We thank Janice Britton-Davidian, Josette Catalan and Marco Perriat-Sanguinet for their help in dissecting mice. We are very also grateful to Leslie Turner and Bettina Harr for sharing with us candidate hybrid sterility region genomic coordinates.

## Data, scripts, code, and supplementary information availability

Raw mapping files (BAM files) and metadata have been uploaded to the Short Read Archive (SRA, NCBI) under the bioproject ID PRJNA603262 (https://www.ncbi.nlm.nih.gov/bioproject/PRJNA603262/).

Custom scripts produced for data analysis are deposited in Zenodo (https://doi.org/10.5281/zenodo.7225771).

Supplementary information is deposited in Zenodo (https://doi.org/10.5281/zenodo.6900875). The following supplementary information is available for this article:

Table S1: sample description

Table S2: sequencing and mapping statistics

Table S3: list of BayPass outlier SNPs

Table S4: gene-based results

Table S5: complete list of outlier genes

Table S6: Variant Effect Predictor results

Table S7: DAVID functional enrichment results

Table S8: Comparison of genetic diversity and recombination rate between outlier and non-outlier gene regions

Table S9 : list of annotated genes with significant selective sweep windows (pool-hmm)

Table S10 : significant local score regions

Table S11 : list of outlier gene sets significantly clustered in the genome (called gene clusters)

Figure S1: Distributions of SNP-based *C_2_* values in the Focal and Control tests

Figure S2: Results of permutation tests assessing co-location of outlier genes with coldspots of recombination

Figure S3: Results of permutation tests assessing co-location of outlier genes with candidate hybrid sterility regions

Figure S4: Autocorrelation and partial autocorrelation estimates in genetic differentiation

Figure S5: Local score distribution along the autosomes and the X chromosome

Figure S6: Results of a parameter-free test for homogeneous distribution of outlier genes in the genome

Figure S7: Details of Figure 3 for the three main gene clusters on chromosomes 2, 7 and 9

## Authors’ contributions

CMS: conceptualization, funding acquisition, data analyses, study and data analyses supervision, drafting the article; EL: bioinformatic developments and data analyses; PC: sampling and DNA sample preparation; DS: library preparation and sequencing; MG: BayPass analyses; GG: participation to research formulation, funding acquisition, sampling. All authors contributed to revising and editing the manuscript.

## References

Albrechtova J, Albrecht T, Baird SJE, Macholán M, Rudolfsen G, Munclinger P, Tucker PK, Piálek J. 2012. Sperm-related phenotypes implicated in both maintenance and breakdown of a natural species barrier in the house mouse. Proc. R. Soc. B:rspb20121802. https://doi.org/10.1098/rspb.2012.1802

Anholt RRH. 2020. Evolution of Epistatic Networks and the Genetic Basis of Innate Behaviors. Trends Genet. 36:24–29. https://doi.org/10.1016/j.tig.2019.10.005

Arguello JR, Benton R. 2017. Open questions: Tackling Darwin’s “instincts”: the genetic basis of behavioral evolution. BMC Biol. 15:26. https://doi.org/10.1186/s12915-017-0369-3

Arya GH, Magwire MM, Huang W, Serrano-Negron YL, Mackay TFC, Anholt RRH. 2015. The genetic basis for variation in olfactory behavior in Drosophila melanogaster. Chem. Senses 40:233–243. https://doi.org/10.1093/chemse/bjv001

Auer TO, Khallaf MA, Silbering AF, Zappia G, Ellis K, Álvarez-Ocaña R, Arguello JR, Hansson BS, Jefferis GSXE, Caron SJC, et al. 2020. Olfactory receptor and circuit evolution promote host specialization. Nature 579:402–408. https://doi.org/10.1038/s41586-020-2073-7

Baird SJE, Petružela J, Jaroň I, Škrabánek P, Martínková N. 2022. Genome polarisation for detecting barriers to geneflow. bioRxiv:2022.03.24.485605. https://doi.org/10.1101/2022.03.24.485605

Banker SE, Bonhomme F, Nachman MW. 2022. Bidirectional Introgression between Mus musculus domesticus and Mus spretus. Genome Biol. Evol. 14:evab288. https://doi.org/10.1093/gbe/evab288

Bastide H, Betancourt A, Nolte V, Tobler R, Stöbe P, Futschik A, Schlötterer C. 2013. A Genome-Wide, Fine-Scale Map of Natural Pigmentation Variation in Drosophila melanogaster. PLoS Genet. 9:e1003534. https://doi.org/10.1371/journal.pgen.1003534

Bay RA, Arnegard ME, Conte GL, Best J, Bedford NL, McCann SR, Dubin ME, Chan YF, Jones FC, Kingsley DM, et al. 2017. Genetic Coupling of Female Mate Choice with Polygenic Ecological Divergence Facilitates Stickleback Speciation. Curr. Biol. 27:3344–3349.e4. https://doi.org/10.1016/j.cub.2017.09.037

Bear DM, Lassance J-M, Hoekstra HE, Datta SR. 2016. The Evolving Neural and Genetic Architecture of Vertebrate Olfaction. Curr. Biol. 26:R1039–R1049. https://doi.org/10.1016/j.cub.2016.09.011

Benton R. 2015. Multigene Family Evolution: Perspectives from Insect Chemoreceptors. Trends Ecol. Evol. 30:590–600. https://doi.org/10.1016/j.tree.2015.07.009

Bimova B, Macholan M, Baird SJE, Munclinger P, Dufkova P, Laukaitis CM, Karn RC, Luzynski K, Tucker PK, Pialek J. 2011. Reinforcement selection acting on the European house mouse hybrid zone. Mol. Ecol. 20:2403–2424. https://doi.org/10.1111/j.1365-294X.2011.05106.x

Boitard S, Kofler R, Françoise P, Robelin D, Schlötterer C, Futschik A. 2013. Pool-hmm: a Python program for estimating the allele frequency spectrum and detecting selective sweeps from next generation sequencing of pooled samples. Mol. Ecol. Resour. 13:337–340. https://doi.org/10.1111/1755-0998.12063

Bolger AM, Lohse M, Usadel B. 2014. Trimmomatic: a flexible trimmer for Illumina sequence data. Bioinformatics 30:2114–2120. https://doi.org/10.1093/bioinformatics/btu170

Boursot P, Din W, Anand R, Darviche D, Dod B, Von Deimling F, Talwar GP, Bonhomme F. 1996. Origin and radiation of the house mouse: mitochondrial DNA phylogeny. J. Evol. Biol. 9:391–415. https://doi.org/10.1046/j.1420-9101.1996.9040391.x

Britton-Davidan J, Fel-Clair F, Lopez J, Alibert P, Boursot P. 2005. Postzygotic isolation between the two European subspecies of the house mouse: estimates from fertility patterns in wild and laboratory-bred hybrids. Biol. J. Linn. Soc. 84:379–393. https://doi.org/10.1111/j.1095-8312.2005.00441.x

Butlin RK, Smadja CM. 2018. Coupling, reinforcement, and speciation. Am. Nat. 191:155–172. https://doi.org/10.1086/695136

Chamero P, Leinders-Zufall T, Zufall F. 2012. From genes to social communication: molecular sensing by the vomeronasal organ. Trends Neurosci. 35:597–606. https://doi.org/10.1016/j.tins.2012.04.011

Chung J, Jun GR, Dupuis J, Farrer LA. 2019. Comparison of methods for multivariate gene-based association tests for complex diseases using common variants. Eur. J. Hum. Genet. 27:811–823. https://doi.org/10.1038/s41431-018-0327-8

Clemente F, Gautier M, Vitalis R. 2018. Inferring sex-specific demographic history from SNP data. PLoS Genet. 14:e1007191.https://doi.org/10.1371/journal.pgen.1007191

Cruickshank TE, Hahn MW. 2014. Reanalysis suggests that genomic islands of speciation are due to reduced diversity, not reduced gene flow. Mol. Ecol. 23:3133–3157. https://doi.org/10.1111/mec.12796

Dickson LB, Merkling SH, Gautier M, Ghozlane A, Jiolle D, Paupy C, Ayala D, Moltini-Conclois I, Fontaine A, Lambrechts L. 2020. Exome-wide association study reveals largely distinct gene sets underlying specific resistance to dengue virus types 1 and 3 in *Aedes aegypti*. PLOS Genet. 16:e1008794. https://doi.org/10.1371/journal.pgen.1008794

Fagny M, Austerlitz F. 2021. Polygenic Adaptation: Integrating Population Genetics and Gene Regulatory Networks. Trends Genet. 37:631–638. https://doi.org/10.1016/j.tig.2021.03.005

Fariello MI, Boitard S, Mercier S, Robelin D, Faraut T, Arnould C, Recoquillay J, Bouchez O, Salin G, Dehais P, et al. 2017. Accounting for Linkage Disequilibrium in genome scans for selection without individual genotypes: the local score approach. Mol. Ecol. 26:3700–3714. https://doi.org/10.1111/mec.14141

Felsenstein J. 1981. Skepticism towards Santa Rosali, or why are there so few kinds of animals? Evolution (N. Y). 35:124–138. https://doi.org/10.1111/j.1558-5646.1981.tb04864.x

Ferretti L, Ramos-Onsins SE, Pérez-Enciso M. 2013. Population genomics from pool sequencing. Mol. Ecol. 22:5561–5576. https://doi.org/10.1111/mec.12522

Fustier M-AM-A., Brandenburg J-T. J-T, Boitard S, Lapeyronnie J, Eguiarte LE, Vigouroux Y, Manicacci D, Tenaillon MI. 2017. Signatures of local adaptation in lowland and highland teosintes from whole-genome sequencing of pooled samples. Mol. Ecol. 26:2738–2756. https://doi.org/10.1111/mec.14082

Futschik A, Schlotterer C. 2010. The Next Generation of Molecular Markers From Massively Parallel Sequencing of Pooled DNA Samples. Genetics 186:207–218. https://doi.org/10.1534/genetics.110.114397

Ganem G, Litel C, Lenormand T. 2008. Variation in mate preference across a house mouse hybrid zone. Heredity (Edinb). 100:594–601. https://doi.org/10.1038/hdy.2008.20

Garner AG, Goulet BE, Farnitano MC, Molina-Henao YF, Hopkins R. 2018. Genomic Signatures of Reinforcement. Genes (Basel). 9. https://doi.org/10.3390/genes9040191

Gautier M. 2015. Genome-Wide Scan for Adaptive Divergence and Association with Population-Specific Covariates. Genetics 201:1555–1579. https://doi.org/10.1534/genetics.115.181453

Gautier M, Foucaud J, Gharbi K, Cézard T, Galan M, Loiseau A, Thomson M, Pudlo P, Kerdelhué C, Estoup A. 2013. Estimation of population allele frequencies from next-generation sequencing data: pool-versus individual-based genotyping. Mol. Ecol. 22:3766–3779. https://doi.org/10.1111/mec.12360

Gautier M, Vitalis R, Flori L, Estoup A. 2022. *f*-Statistics estimation and admixture graph construction with Pool-Seq or allele count data using the R package poolfstat. Mol. Ecol. Resour. 22:1394–1416. https://doi.org/10.1111/1755-0998.13557

Gel B, Díez-Villanueva A, Serra E, Buschbeck M, Peinado MA, Malinverni R. 2015. regioneR: an R/Bioconductor package for the association analysis of genomic regions based on permutation tests. Bioinformatics 32:btv562.https://doi.org/10.1093/bioinformatics/btv562

Geraldes A, Basset P, Smith KL, Nachman MW. 2011. Higher differentiation among subspecies of the house mouse (Mus musculus) in genomic regions with low recombination. Mol. Ecol. 20:4722–4736. https://doi.org/10.1111/j.1365-294X.2011.05285.x

Gu Z, Eils R, Schlesner M. 2016. Gtrellis: An R/Bioconductor package for making genome-level Trellis graphics. BMC Bioinformatics 17:169. https://doi.org/10.1186/s12859-016-1051-4

Haasl RJ, Payseur BA. 2016. Fifteen years of genome-wide scans for selection: trends, lessons, and unaddressed genetic sources of complication. Mol. Ecol. 25:5-23 https://doi.org/10.1111/mec.13339

Hahn MW. 2006. Accurate Inference and Estimation in Population Genomics. Mol. Biol. Evol. 23:911–918. https://doi.org/10.1093/molbev/msj094

Haque MM, Nilsson EE, Holder LB, Skinner MK. 2016. Genomic Clustering of differential DNA methylated regions (epimutations) associated with the epigenetic transgenerational inheritance of disease and phenotypic variation. BMC Genomics 17:418. https://doi.org/10.1186/s12864-016-2748-5

Hayden S, Bekaert M, Crider TA, Mariani S, Murphy WJ, Teeling EC. 2010. Ecological adaptation determines functional mammalian olfactory subgenomes. Genome Res. 20:1–9. https://doi.org/10.1101/gr.099416.109

Hayden S, Bekaert M, Goodbla A, Murphy WJ, Dávalos LM, Teeling EC. 2014. A cluster of olfactory receptor genes linked to frugivory in bats. Mol. Biol. Evol. 31:917–927. https://doi.org/10.1093/molbev/msu043

Hayden S, Teeling EC. 2014. The molecular biology of vertebrate olfaction. Anat. Rec. (Hoboken). 297:2216–2226. https://doi.org/10.1002/ar.23031

Hench K, Vargas M, Höppner MP, McMillan WO, Puebla O. 2019. Inter-chromosomal coupling between vision and pigmentation genes during genomic divergence. Nat. Ecol. Evol. 3:657–667. https://doi.org/10.1038/s41559-019-0814-5

Hivert V, Leblois R, Petit EJ, Gautier M, Vitalis R. 2018. Measuring Genetic Differentiation from Pool-seq Data. Genetics 210:315–330. https://doi.org/10.1534/genetics.118.300900

Hohenlohe PA, Phillips PC, Cresko WA. 2010. Using population genomics to detect selection in natural populations: key concepts and methodological considerations. Int. J. Plant Sci. 171:1059–1071. https://doi.org/10.1086/656306

Hopkins R, Rausher MD. 2011. Identification of two genes causing reinforcement in the Texas wildflower Phlox drummondii. Nature 469:411. https://doi.org/10.1038/nature09641

Huang DW, Sherman BT, Lempicki RA. 2009. Systematic and integrative analysis of large gene lists using DAVID bioinformatics resources. Nat. Protoc. 4:44–57. https://doi.org/10.1038/nprot.2008.211

Huang H, Chanda P, Alonso A, Bader JS, Arking DE. 2011. Gene-Based Tests of Association. PLoS Genet. 7:e1002177. https://doi.org/10.1371/journal.pgen.1002177

Hurst JL, Beynon RJ, Armstrong SD, Davidson AJ, Roberts SA, Gómez-Baena G, Smadja CM, Ganem G. 2017. Molecular heterogeneity in major urinary proteins of *Mus musculus* subspecies: potential candidates involved in speciation. Sci. Rep. 7:44992. https://doi.org/10.1038/srep44992

Ibarra-Soria X, Levitin M, Logan D. 2014. The genomic basis of vomeronasal-mediated behaviour. Mamm. Genome 25:75–86. https://doi.org/10.1007/s00335-013-9463-1

Ibarra-Soria X, Levitin MO, Saraiva LR, Logan DW. 2014. The Olfactory Transcriptomes of Mice. PLoS Genet. 10:e1004593. https://doi.org/10.1371/journal.pgen.1004593

Isogai Y, Si S, Pont-Lezica L, Tan T, Kapoor V, Murthy VN, Dulac C. 2011. Molecular organization of vomeronasal chemoreception. Nature 478:241–245. https://doi.org/10.1038/nature10437

Jiao H, Hong W, Nevo E, Li K, Zhao H. 2019. Convergent reduction of V1R genes in subterranean rodents. BMC Evol. Biol. 19:176. https://doi.org/10.1186/s12862-019-1502-4

Jourjine N, Hoekstra HE. 2021. Expanding evolutionary neuroscience: insights from comparing variation in behavior. Neuron 109:1084–1099. https://doi.org/10.1016/j.neuron.2021.02.002

Kent CF, Tiwari T, Rose S, Patel H, Conflitti IM, Zayed A. 2019. Studying the genetics of behavior in the genomics era. In: Encyclopedia of Animal Behavior. Elsevier. p. 223–233. https://doi.org/10.1016/B978-0-12-809633-8.90054-2

Kern AD, Schrider DR. 2018. diploS/HIC: An Updated Approach to Classifying Selective Sweeps. G3 Genes|Genomes|Genetics 8:1959–1970. https://doi.org/10.1534/g3.118.200262

Khan I, Yang Z, Maldonado E, Li C, Zhang G, Gilbert MTP, Jarvis ED, O’Brien SJ, Johnson WE, Antunes A. 2015. Olfactory Receptor Subgenomes Linked with Broad Ecological Adaptations in Sauropsida. Mol. Biol. Evol. 32:2832–2843. https://doi.org/10.1093/molbev/msv155

Kirkpatrick M, Servedio MR. 1999. The reinforcement of mating preferences on an island. Genetics 151:865–884. https://doi.org/10.1093/genetics/151.2.865

Kofler R, Pandey R V, Schlotterer C. 2011. PoPoolation2: identifying differentiation between populations using sequencing of pooled DNA samples (Pool-Seq). Bioinformatics 27:3435–3436. https://doi.org/10.1093/bioinformatics/btr589

Kopp M, Servedio MR, Mendelson TC, Safran RJ, Rodríguez RL, Hauber ME, Scordato EC, Symes LB, Balakrishnan CN, Zonana DM, et al. 2018. Mechanisms of Assortative Mating in Speciation with Gene Flow: Connecting Theory and Empirical Research. Am. Nat. 191:1–20. https://doi.org/10.1086/694889

Langmead B, Salzberg SL. 2012. Fast gapped-read alignment with Bowtie 2. Nat. Methods 9:357–359. https://doi.org/10.1038/nmeth.1923

Latour Y, Perriat-Sanguinet M, Caminade P, Boursot P, Smadja CMCM, Ganem G. 2014. Sexual selection against natural hybrids may contribute to reinforcement in a house mouse hybrid zone. Proc. R. Soc. London B Biol. Sci. 281. https://doi.org/10.1098/rspb.2013.2733

Latta RG. 1998. Differentiation of allelic frequencies at quantitative trait loci affecting locally adaptive traits. Am. Nat. 151:283–292. https://doi.org/10.1086/286119

Laturney M, Moehring AJ. 2012. The Genetic Basis of Female Mate Preference and Species Isolation in Drosophila. Int. J. Evol. Biol. 2012:1–13. https://doi.org/10.1155/2012/328392

Laurie CC, Nickerson DA, Anderson AD, Weir BS, Livingston RJ, Dean MD, Smith KL, Schadt EE, Nachman MW. 2007. Linkage disequilibrium in wild mice. Plos Genet. 3:1487–1495. https://doi.org/10.1371/journal.pgen.0030144

Li H, Handsaker B, Wysoker A, Fennell T, Ruan J, Homer N, Marth G, Abecasis G, Durbin R, 1000 Genome Project Data Processing Subgroup. 2009. The Sequence Alignment/Map format and SAMtools. Bioinformatics 25:2078–2079. https://doi.org/10.1093/bioinformatics/btp352

Li K, Hong W, Jiao H, Wang G-D, Rodriguez KA, Buffenstein R, Zhao Y, Nevo E, Zhao H. 2015. Sympatric speciation revealed by genome-wide divergence in the blind mole rat Spalax. Proc. Natl. Acad. Sci. U. S. A. 112:11905–11910. https://doi.org/10.1073/pnas.1514896112

Liberles SD. 2014. Mammalian Pheromones. Annu. Rev. Physiol. 76:151–175. https://doi.org/10.1146/annurev-physiol-021113-170334

Librado P, Rozas J. 2016. Weak polygenic selection drives the rapid adaptation of the chemosensory system: Lessons from the upstream regions of the major gene families. Genome Biol. Evol. 8:2493–2504. https://doi.org/10.1093/gbe/evw191

Loire E, Tusso S, Caminade P, Severac D, Boursot P, Ganem G, Smadja CM. 2017. Do changes in gene expression contribute to sexual isolation and reinforcement in the house mouse? Mol. Ecol. 26:5189–5202.https://doi.org/10.1111/mec.14212

McLaren W, Gil L, Hunt SE, Riat HS, Ritchie GRS, Thormann A, Flicek P, Cunningham F. 2016. The Ensembl Variant Effect Predictor. Genome Biol. 17:122. https://doi.org/10.1186/s13059-016-0974-4

Meier JI, Salazar PA, Kučka M, Davies RW, Dréau A, Aldás I, Power OB, Nadeau NJ, Bridle JR, Rolian C, et al. 2021. Haplotype tagging reveals parallel formation of hybrid races in two butterfly species. Proc. Natl. Acad. Sci. U. S. A. 118. https://doi.org/10.1073/pnas.2015005118

Mendelson TC, Safran RJ. 2021. Speciation by sexual selection: 20 years of progress. Trends Ecol. Evol. 36:1153–1163. https://doi.org/10.1016/j.tree.2021.09.004

Merrill RM, Rastas P, Martin SH, Melo MC, Barker S, Davey J, McMillan WO, Jiggins CD. 2019. Genetic dissection of assortative mating behavior. PLOS Biol. 17:e2005902. https://doi.org/10.1371/journal.pbio.2005902

Morgan AP, Gatti DM, Najarian ML, Keane TM, Galante RJ, Pack AI, Mott R, Churchill GA, de Villena FP-M. 2017. Structural Variation Shapes the Landscape of Recombination in Mouse. Genetics 206:603–619. https://doi.org/10.1534/genetics.116.197988

Mukaj A, Piálek J, Fotopulosova V, Morgan AP, Odenthal-Hesse L, Parvanov ED, Forejt J. 2020. Prdm9 inter-subspecific interactions in hybrid male sterility of house mouse. Mol. Biol. Evol. 37:3423–3438. https://doi.org/10.1093/molbev/msaa167

Neale BM, Sham PC. 2004. The future of association studies: gene-based analysis and replication. Am. J. Hum. Genet. 75:353–362. https://doi.org/10.1086/423901

Niepoth N, Bendesky A. 2020. How Natural Genetic Variation Shapes Behavior. Annu. Rev. Genomics Hum. Genet. 21:437–463. https://doi.org/10.1146/annurev-genom-111219-080427

North H, Caminade P, Severac D, Belkhir K, Smadja C. 2020. The role of copy-number variation in the reinforcement of sexual isolation between the two European subspecies of the house mouse. Philos. Trans. R. Soc. B Biol. Sci. 375:8 https://doi.org/10.1098/rstb.2019.0540

Olazcuaga L, Loiseau A, Parrinello H, Paris M, Fraimout A, Guedot C, Diepenbrock LM, Kenis M, Zhang J, Chen X, et al. 2021. A whole-genome scan for association with invasion success in the fruit fly *Drosophila suzukii* using contrasts of allele frequencies corrected for population structure. Mol. Biol. Evol. 37:2369–2385. https://doi.org/10.1093/molbev/msaa098

Ortiz-Barrientos D, Counterman BA, Noor MAF. 2004. The genetics of speciation by reinforcement. Plos Biol. 2:e416.https://doi.org/10.1371/journal.pbio.0020416

Ospina OE, Lemmon AR, Dye M, Zdyrski C, Holland S, Stribling D, Kortyna ML, Lemmon EM. 2021. Neurogenomic divergence during speciation by reinforcement of mating behaviors in chorus frogs (Pseudacris). BMC Genomics 22:711.https://doi.org/10.1186/s12864-021-07995-3

Pardy JA, Lahib S, Noor MAF, Moehring AJ. 2021. Intraspecific genetic variation for behavioral isolation loci in drosophila. Genes (Basel). 12. https://doi.org/10.3390/genes12111703

Paudel Y, Madsen O, Megens H-J, Frantz LAF, Bosse M, Crooijmans RPMA, Groenen MAM. 2015. Copy number variation in the speciation of pigs: a possible prominent role for olfactory receptors. BMC Genomics 16:330. https://doi.org/10.1186/s12864-015-1449-9

Poelstra JW, Richards EJ, Martin CH. 2018. Speciation in sympatry with ongoing secondary gene flow and a potential olfactory trigger in a radiation of Cameroon cichlids. Mol. Ecol. 27:4270–4288. https://doi.org/10.1111/mec.14784

Pritchard JK, Pickrell JK, Coop G. 2010. The genetics of human adaptation: hard sweeps, soft sweeps, and polygenic adaptation. Curr. Biol. 20:R208–15. https://doi.org/10.1016/j.cub.2009.11.055

Quinlan AR, Hall IM. 2010. BEDTools: a flexible suite of utilities for comparing genomic features. Bioinformatics 26:841–842. https://doi.org/10.1093/bioinformatics/btq033

R Core Team. 2019. R: A Language and Environment for Statistical Computing. Available online at https://www.R-project.org/

Raufaste N, Orth A, Belkhir K, Senet D, Smadja C, Baird SJEJE, Bonhomme F, Dod B, Boursot P. 2005. Inference of selection and migration in the danish house mouse hybrid zone. Biol. J. Linn. Soc. 84:593–616. https://doi.org/10.1111/j.1095-8312.2005.00457.x

Ravinet M, Faria R, Butlin RK, Galindo J, Bierne N, Rafajlović M, Noor MAF, Mehlig B, Westram AM. 2017. Interpreting the genomic landscape of speciation: a road map for finding barriers to gene flow. J. Evol. Biol. 30:1450–1477. https://doi.org/10.1111/jeb.13047

Roda F, Walter GM, Nipper R, Ortiz-Barrientos D. 2017. Genomic clustering of adaptive loci during parallel evolution of an Australian wildflower. Mol. Ecol. 26:3687–3699. https://doi.org/10.1111/mec.14150

Roesti M, Gavrilets S, Hendry AP, Salzburger W, Berner D. 2014. The genomic signature of parallel adaptation from shared genetic variation. Mol. Ecol. 23:3944–3956. https://doi.org/10.1111/mec.12720

Sánchez-Gracia A, Vieira FG, Almeida FC, Rozas J. 2011. Comparative Genomics of the Major Chemosensory Gene Families in Arthropods. In: eLS. John Wiley & Sons, Ltd. https://doi.org/10.1002/9780470015902.a0022848

Santos PSC, Mezger M, Kolar M, Michler F-U, Sommer S. 2018. The best smellers make the best choosers: mate choice is affected by female chemosensory receptor gene diversity in a mammal. Proc. R. Soc. B Biol. Sci. 285:20182426.https://doi.org/10.1098/rspb.2018.2426

Schlötterer C, Tobler R, Kofler R, Nolte V. 2014. Sequencing pools of individuals – mining genome-wide polymorphism data without big funding. Nat. Rev. Genet. 15:749–763. https://doi.org/10.1038/nrg3803

Schneider K, White TJ, Mitchell S, Adams CE, Reeve R, Elmer KR. 2021. The pitfalls and virtues of population genetic summary statistics: Detecting selective sweeps in recent divergences. J. Evol. Biol. 34:893–909. https://doi.org/10.1111/jeb.13738

Sella G, Barton NH. 2019. Thinking About the Evolution of Complex Traits in the Era of Genome-Wide Association Studies. Annu. Rev. Genomics Hum. Genet. 20:461–493. https://doi.org/10.1146/annurev-genom-083115-022316

Servedio MR. 2009. The role of linkage disequilibrium in the evolution of premating isolation. Heredity (Edinb). 102:51–56. https://doi.org/10.1038/hdy.2008.98

Servedio MR, Boughman JW. 2017. The Role of Sexual Selection in Local Adaptation and Speciation. Annu. Rev. Ecol. Evol. Syst. 48:85–109. https://doi.org/10.1146/annurev-ecolsys-110316-022905

Servedio MR, Noor MAF. 2003. The role of reinforcement in speciation: Theory and Data. Annu. Rev. Ecol. Syst. 34:339–364. https://doi.org/10.1146/annurev.ecolsys.34.011802.132412

Shi P, Zhang JZ. 2007. Comparative genomic analysis identifies an evolutionary shift of vomeronasal receptor gene repertoires in the vertebrate transition from water to land. Genome Res. 17:166–174. https://doi.org/10.1101/gr.6040007

Smadja C, Butlin RK. 2009. On the scent of speciation: the chemosensory system and its role in premating isolation. Heredity (Edinb). 102:77–97. https://doi.org/10.1038/hdy.2008.55

Smadja C, Catalan J, Ganem G. 2004. Strong premating divergence in a unimodal hybrid zone between two subspecies of the house mouse. J. Evol. Biol. 17:165–176. https://doi.org/10.1046/j.1420-9101.2003.00647.x

Smadja C, Ganem G. 2002. Subspecies recognition in the house mouse: a study of two populations from the border of a hybrid zone. Behav. Ecol. 13:312–320. https://doi.org/10.1093/beheco/13.3.312

Smadja C, Ganem G. 2005. Asymmetrical reproductive character displacement in the house mouse. J. Evol. Biol. 18:1485–1493. https://doi.org/10.1111/j.1420-9101.2005.00944.x

Smadja C, Ganem G. 2008. Divergence of odorant signals within and between the two European subspecies of the house mouse. Behav. Ecol. 19:223–230. https://doi.org/10.1093/beheco/arm127

Smadja CM, Butlin RK. 2011. A framework for comparing processes of speciation in the presence of gene flow. Mol. Ecol. 20:5123–5140. https://doi.org/10.1111/j.1365-294X.2011.05350.x

Smadja CM, Canbäck B, Vitalis R, Gautier M, Ferrari J, Zhou J-J, Butlin RK. 2012. Large-scale candidate gene scan reveals the role of chemoreceptor genes in host plant specialization and speciation in the pea aphid. Evolution (N. Y). 66:2723–2738. https://doi.org/10.1111/j.1558-5646.2012.01612.x

Smadja CM, Loire E, Caminade P, Thoma M, Latour Y, Roux C, Thoss M, Penn DJDJ, Ganem G, Boursot P. 2015. Seeking signatures of reinforcement at the genetic level: a hitchhiking mapping and candidate gene approach in the house mouse. Mol. Ecol. 24:4222–4237. https://doi.org/10.1111/mec.13301

Smadja CM, Loire E, Caminade P, Severac D, Gautier M, Ganem G. 2022a. Supplementary material of Divergence of olfactory receptors associated with the evolution of assortative mating and reproductive isolation in mice. Zenodo, https://doi.org/10.5281/zenodo.6900875

Smadja CM, Loire E, Caminade P, Severac D, Gautier M, Ganem G. 2022b. Scripts of Divergence of olfactory receptors associated with the evolution of assortative mating and reproductive isolation in mice. Zenodo, https://doi.org/10.5281/zenodo.7225771

Staubach F, Lorenc A, Messer PW, Tang K, Petrov DA, Tautz D. 2012. Genome Patterns of Selection and Introgression of Haplotypes in Natural Populations of the House Mouse (Mus musculus). PLoS Genet 8:e1002891. https://doi.org/10.1371/journal.pgen.1002891

Stölting KN, Paris M, Meier C, Heinze B, Castiglione S, Bartha D, Lexer C. 2015. Genome-wide patterns of differentiation and spatially varying selection between postglacial recolonization lineages of *Populus alba* (Salicaceae), a widespread forest tree. New Phytol. 207:723–734. https://doi.org/10.1111/nph.13392

Swarup S, Huang W, Mackay TFC, Anholt RRH. 2013. Analysis of natural variation reveals neurogenetic networks for *Drosophila* olfactory behavior. Proc. Natl. Acad. Sci. U. S. A. 110:1017–1022. https://doi.org/10.1073/pnas.1220168110

Tajima F. 1983. Evolutionary relationship of DNA sequences in finite populations. Genetics 105:437–460. https://doi.org/10.1093/genetics/105.2.437

Toews DPL, Taylor SA, Streby HM, Kramer GR, Lovette IJ. 2019. Selection on VPS13A linked to migration in a songbird. Proc. Natl. Acad. Sci. U. S. A. 116:18272–18274. https://doi.org/10.1073/pnas.1909186116

Turner LM, Harr B. 2014. Genome-wide mapping in a house mouse hybrid zone reveals hybrid sterility loci and Dobzhansky-Muller interactions. Elife 3:e02504.https://doi.org/10.7554/eLife.02504

Turner LM, Schwahn DJ, Harr B. 2012. Reduced male fertility is common but highly variable in form and severity in a natural jouse mouse hybrid zone. Evolution (N. Y). 66:443–458. https://doi.org/10.1111/j.1558-5646.2011.01445.x

Ullrich KK, Linnenbrink M, Tautz D. 2017. Introgression patterns between house mouse subspecies and species reveal genomic windows of frequent exchange. bioRxiv:168328.https://doi.org/10.1101/168328

Vijay N, Bossu CM, Poelstra JW, Weissensteiner MH, Suh A, Kryukov AP, Wolf Jochen B. W., Wolf J. B. W., Lindell J, Backstrom N, et al. 2016. Evolution of heterogeneous genome differentiation across multiple contact zones in a crow species complex. Nat. Commun. 7:13195.https://doi.org/10.1038/ncomms13195

Wang M, Huang J, Liu Y, Ma L, Potash JB, Han S. 2017. COMBAT: A Combined Association Test for Genes Using Summary Statistics. Genetics 207:883–891. https://doi.org/10.1534/genetics.117.300257

Wang Y, Reddiex AJ, Allen SL, Chenoweth SF. 2022. Direct and indirect impacts of positive selection on genomic variation in Drosophila serrata. bioRxiv 2022.03.31.486660. https://doi.org/10.1101/2022.03.31.486660

Waterston RH, Lindblad-Toh K, Birney E, Rogers J, Abril JF, Agarwal P, Agarwala R, Ainscough R, Alexandersson M, An P, et al. 2002. Initial sequencing and comparative analysis of the mouse genome. Nature 420:520–562. https://doi.org/10.1038/nature01262

Watterson GA. 1975. On the number of segregating sites in genetical models without recombination. Theor. Popul. Biol. 7:256–276. https://doi.org/10.1016/0040-5809(75)90020-9

Wolf JBW, Ellegren H. 2017. Making sense of genomic islands of differentiation in light of speciation. Nat. Rev. Genet. 18:87–100. https://doi.org/10.1038/nrg.2016.133

Yang H, Zhang YP. 2007. Genomic organization and sequence analysis of the vomeronasal receptor V2R genes in mouse genome. Chinese Sci. Bull. 52:336–342. https://doi.org/10.1007/s11434-007-0059-6

Yang J, Jiang H, Yeh C-T, Yu J, Jeddeloh JA, Nettleton D, Schnable PS. 2015. Extreme-phenotype genome-wide association study (XP-GWAS): a method for identifying trait-associated variants by sequencing pools of individuals selected from a diversity panel. Plant J. 84:587–596. https://doi.org/10.1111/tpj.13029

Yeaman S. 2013. Genomic rearrangements and the evolution of clusters of locally adaptive loci. Proc. Natl. Acad. Sci. U. S. A. 110:E1743-E1751. https://doi.org/10.1073/pnas.1219381110

Yeaman S, Whitlock MC. 2011. The genetic architecture of adaptation under migration–selection balance. Evolution (N. Y). 65:1897–1911. https://doi.org/10.1111/j.1558-5646.2011.01269.x

Yohe LR, Leiser-Miller LB, Kaliszewska ZA, Donat P, Santana SE, Dávalos LM. 2021. Diversity in olfactory receptor repertoires is associated with dietary specialization in a genus of frugivorous bat. G3 Genes|Genomes|Genetics 11:jkab260.https://doi.org/10.1093/g3journal/jkab260

Zhang XH, Zhang XM, Firestein S. 2007. Comparative genomics of odorant and pheromone receptor genes in rodents. Genomics 89:441–450. https://doi.org/10.1016/j.ygeno.2007.01.002

Zhang XM, Firestein S. 2002. The olfactory receptor gene superfamily of the mouse. Nat. Neurosci. 5:124–133. https://doi.org/10.1038/nn800

