## Supplementary Figures v2 for "Divergence of olfactory receptors associated with the evolution of assortative mating and reproductive isolation in mice"

**a.**

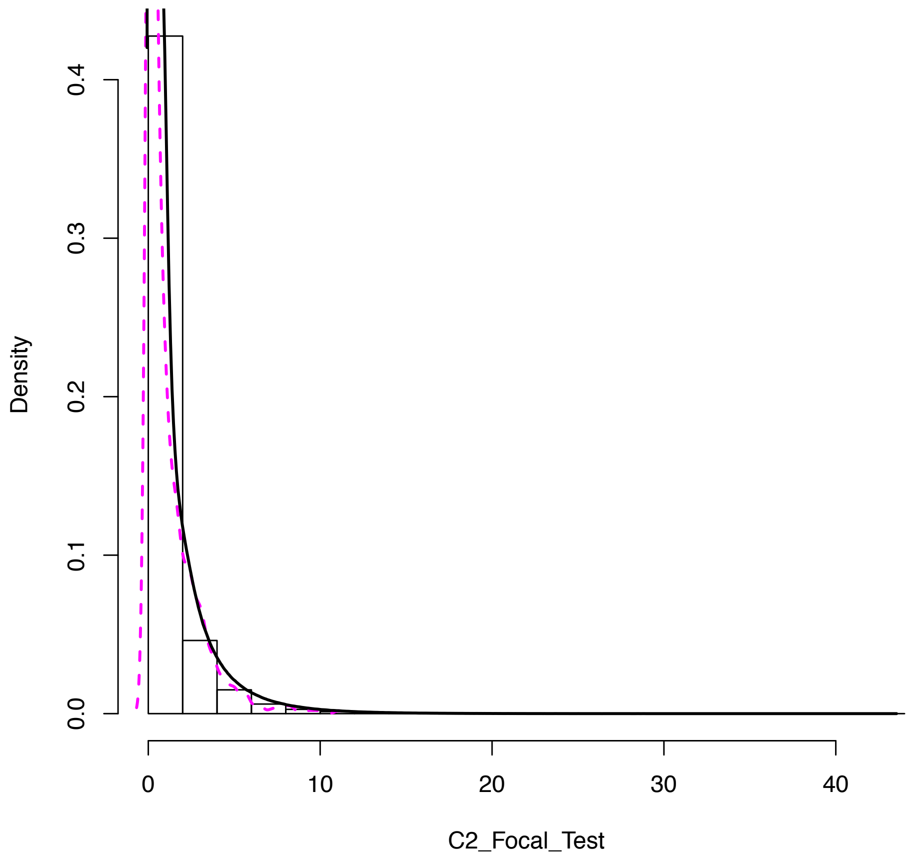

**b.**

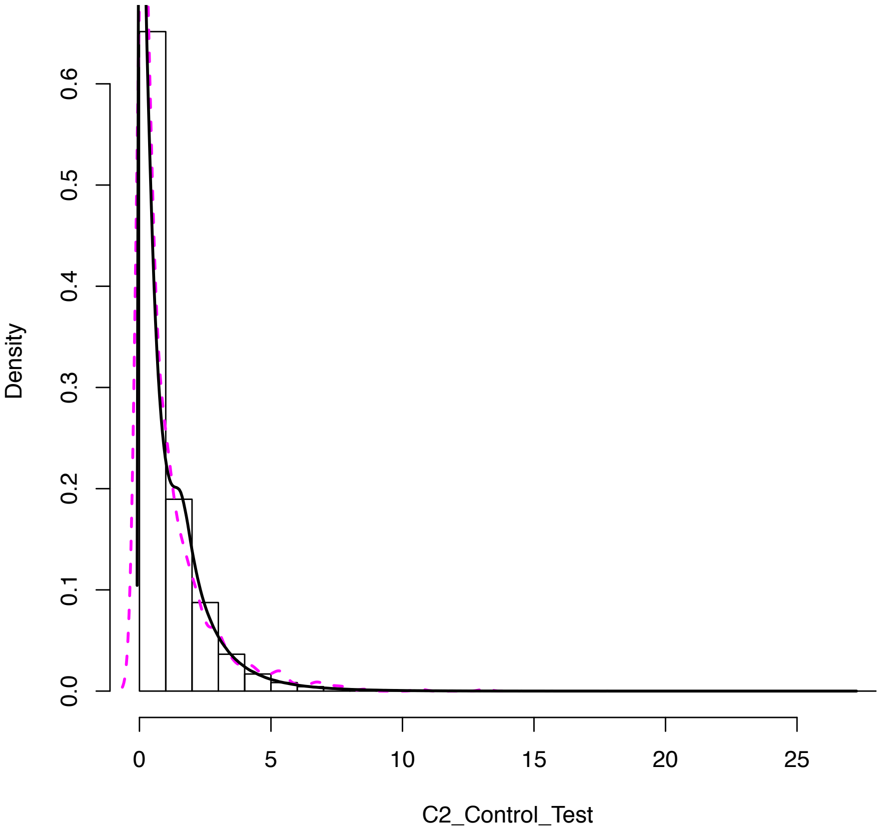


**Supplementary Figure S1** : Distribution of SNP-based $C_2$ values (contrast statistic) computed with the program BAYPASS. a. Focal Test ; b. Control Test. Black lines indicate the empirical probability density functions and magenta dotted lines indicate the expected Chi-squared probability density functions with one degree of freedom.

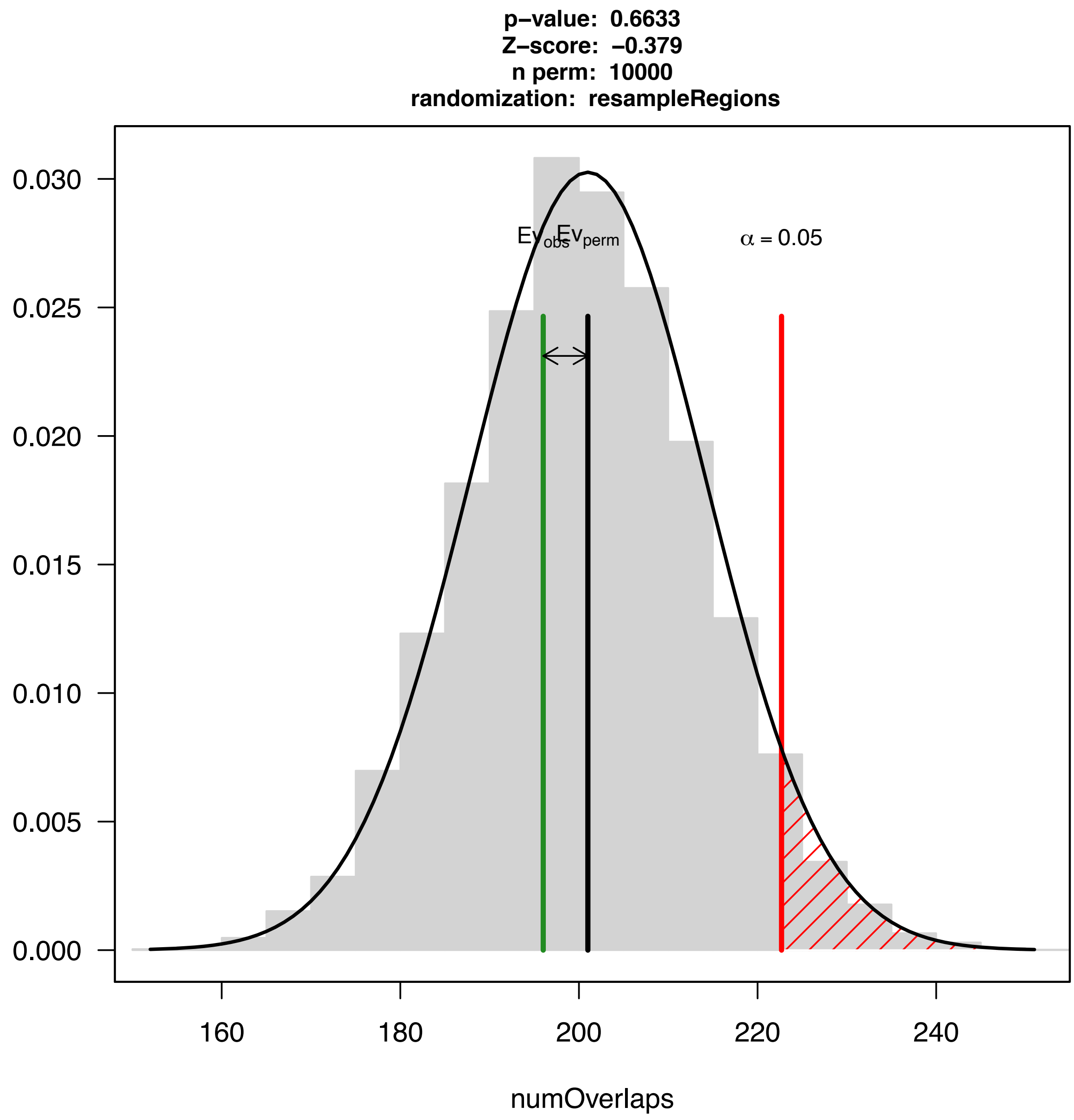


**Supplementary Figure S2**: Results of permutation tests assessing whether outlier genes are overrepresented in coldspots of recombination (as reported by Morgan et al. 2017). 10,000 permutations were conducted.

a. b.

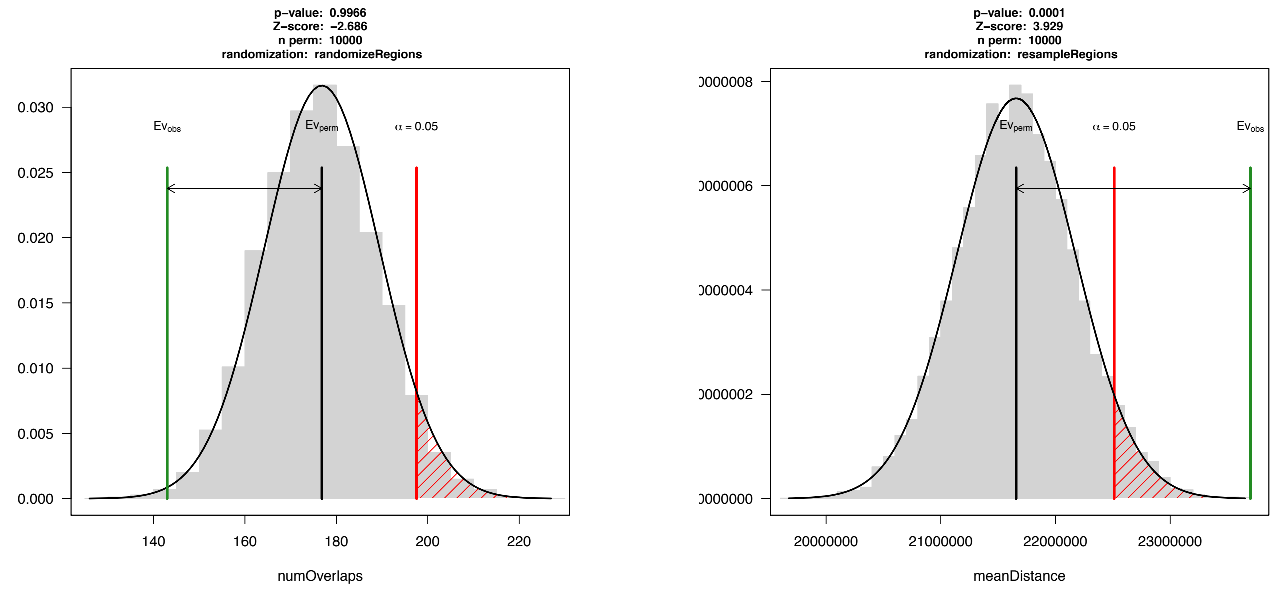


**Supplementary Figure S3**: Results of two permutation tests assessing (a) whether outlier genes are over-represented in genomic regions identified as candidates for hybrid sterility [1,2]; (b) whether outlier genes are closer or more distant to candidate hybrid sterility regions than expected by chance. 10,000 permutations were conducted for each test.

a.

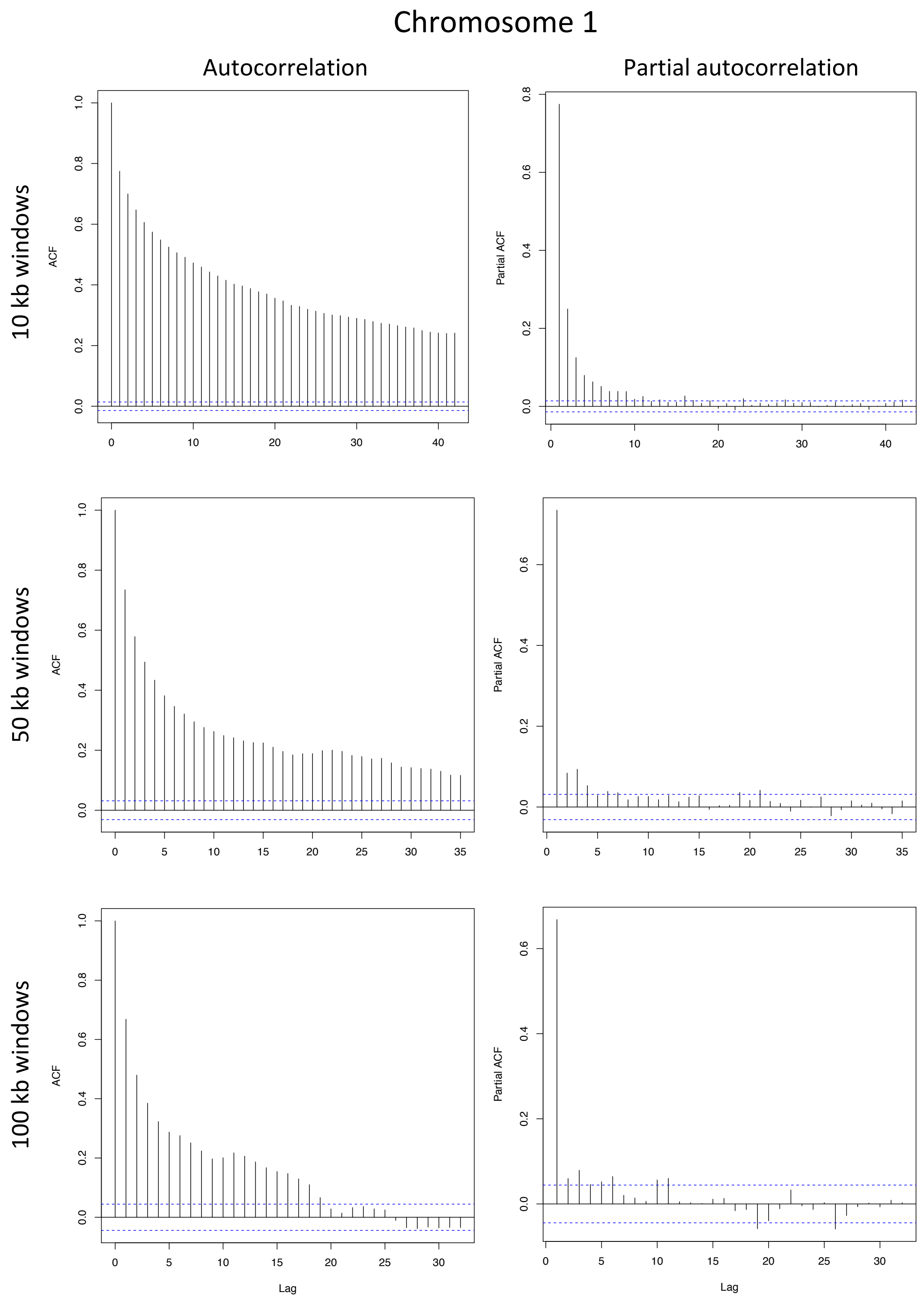
Chromosome 1
Autocorrelation
Partial autocorrelation
10 kb windows
50 kb windows
100 kb windows
ACF
Partial ACF
Lag

b.

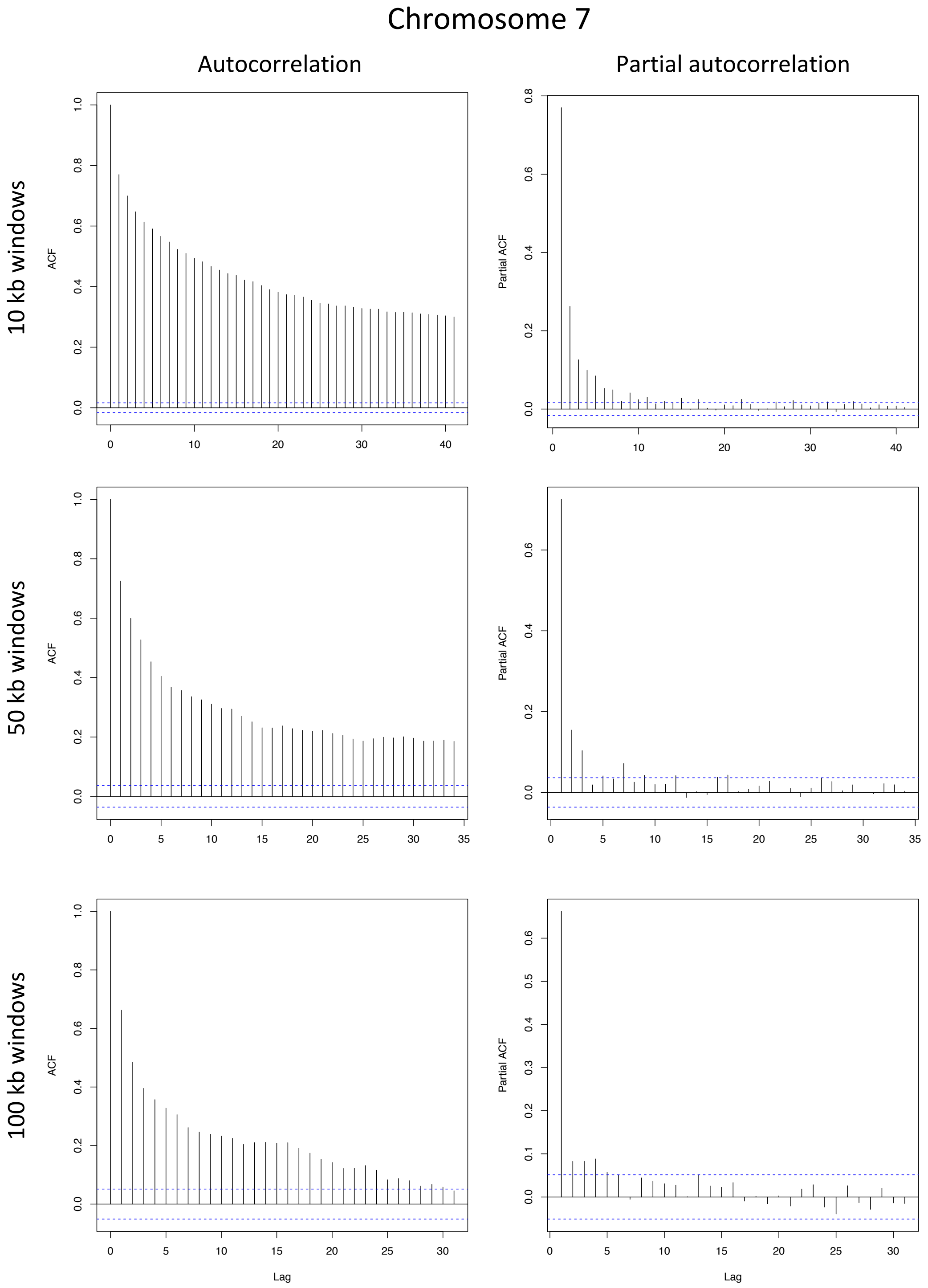
Chromosome 7
Autocorrelation
Partial autocorrelation
10 kb windows
50 kb windows
100 kb windows
ACF
Partial ACF
Lag

c.

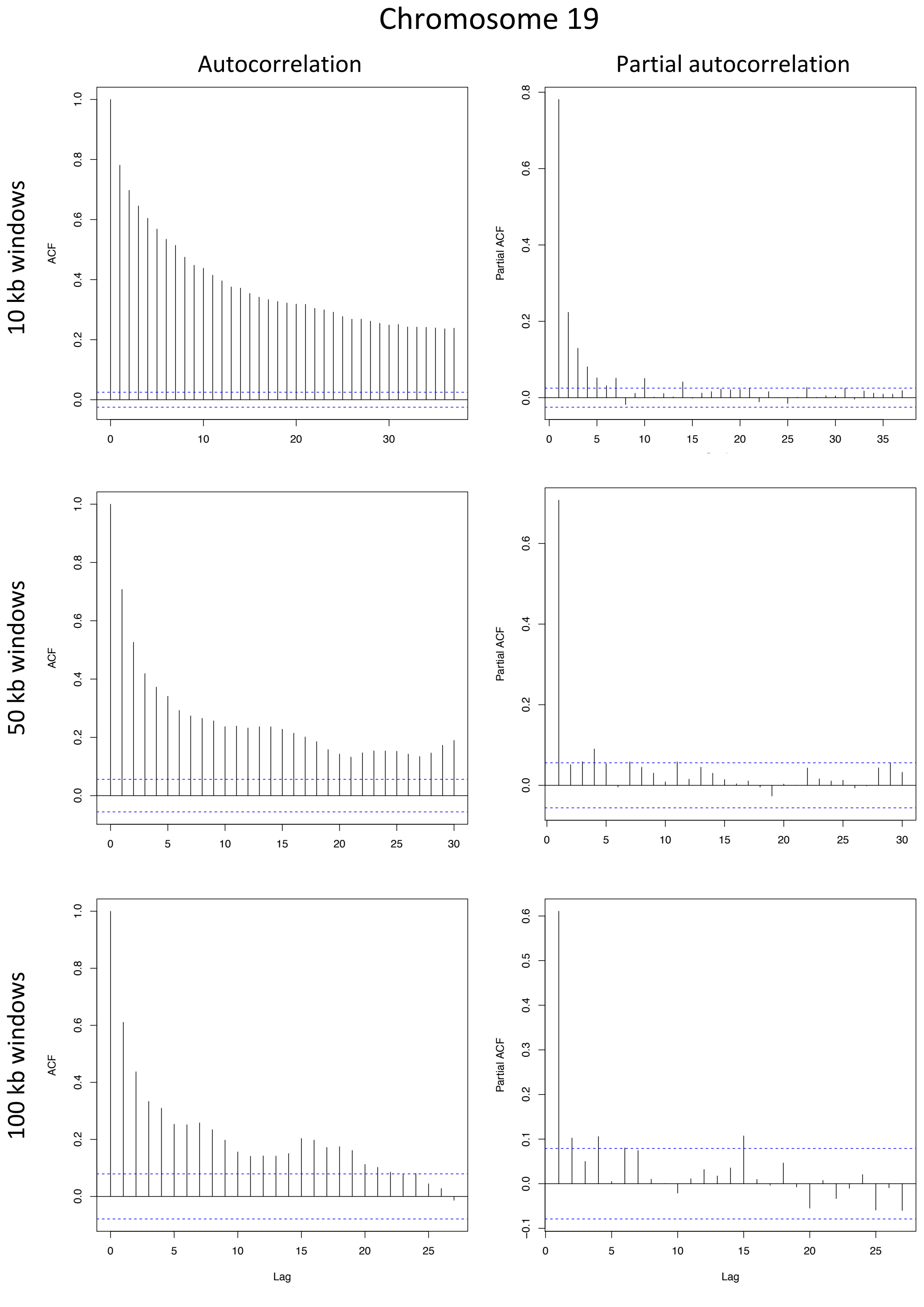


**Supplementary Figure S4** : Autocorrelation (ACF, left panels) and partial autocorrelation (partial ACF, right panels) estimates in genetic differentiation (BAYPASS $C_2$ contrast statistic), measured in 10-kb, 50-kb and 100-kb windows. Results are shown for three chromosomes: (a) chromosome 1 (the longest), (b) chromosome 7 (containing several clusters of outlier genes) and (c) chromosome 19 (the shortest). The auto-correlation at lag 0 is by definition equal to 1. Dashed lines represent the 95% CIs around an autocorrelation coefficient equal to zero.

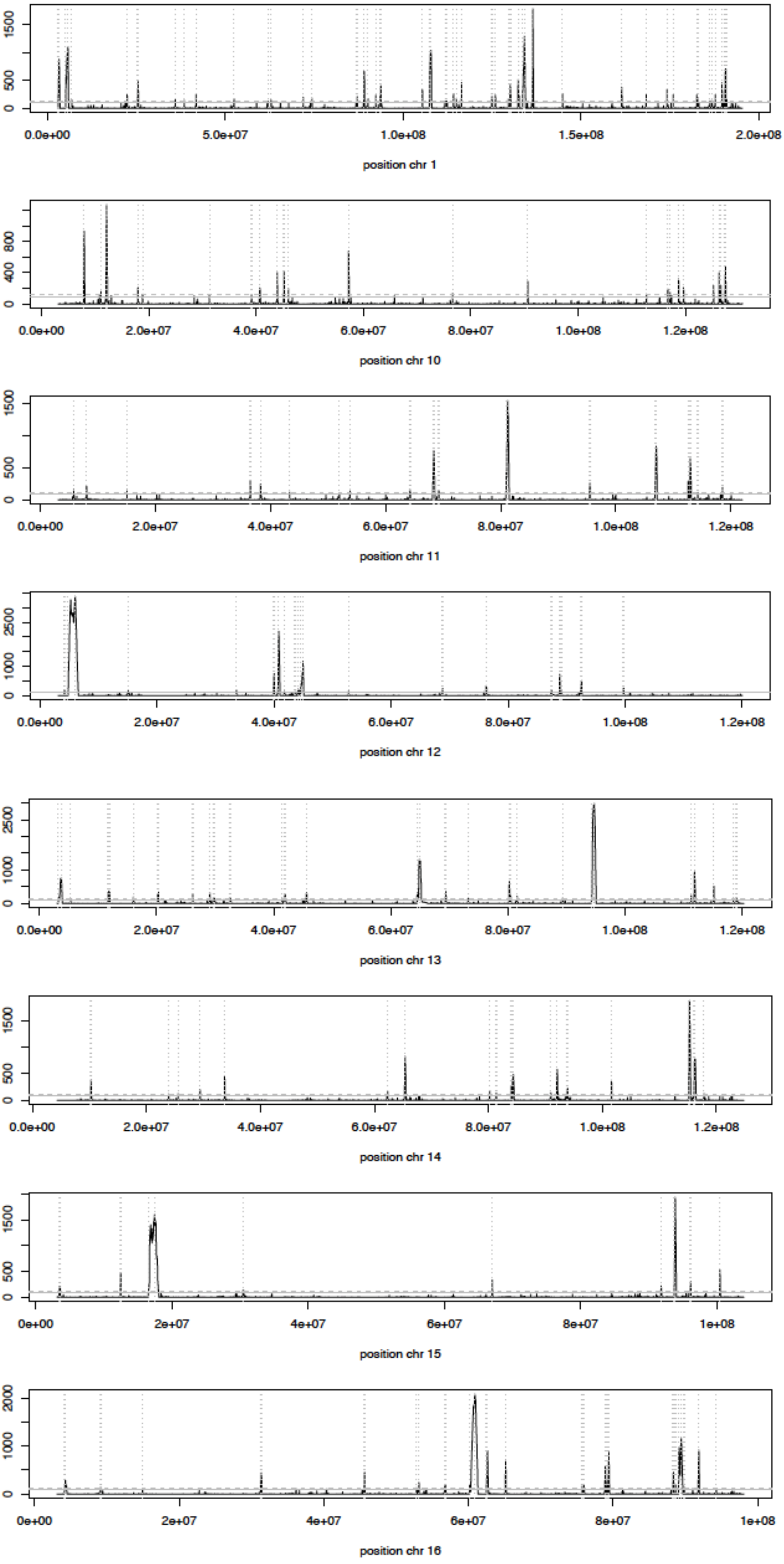
1500
500
0
0.0e+00
5.0e+07
1.0e+08
1.5e+08
2.0e+08
position chr 1
800
400
0
0.0e+00
2.0e+07
4.0e+07
6.0e+07
8.0e+07
1.0e+08
1.2e+08
position chr 10
1500
500
0
0.0e+00
2.0e+07
4.0e+07
6.0e+07
8.0e+07
1.0e+08
1.2e+08
position chr 11
2500
1000
0
0.0e+00
2.0e+07
4.0e+07
6.0e+07
8.0e+07
1.0e+08
1.2e+08
position chr 12
2500
1000
0
0.0e+00
2.0e+07
4.0e+07
6.0e+07
8.0e+07
1.0e+08
1.2e+08
position chr 13
1500
500
0
0.0e+00
2.0e+07
4.0e+07
6.0e+07
8.0e+07
1.0e+08
1.2e+08
position chr 14
1500
500
0
0e+00
2e+07
4e+07
6e+07
8e+07
1e+08
position chr 15
2000
1000
0
0e+00
2e+07
4e+07
6e+07
8e+07
1e+08
position chr 16

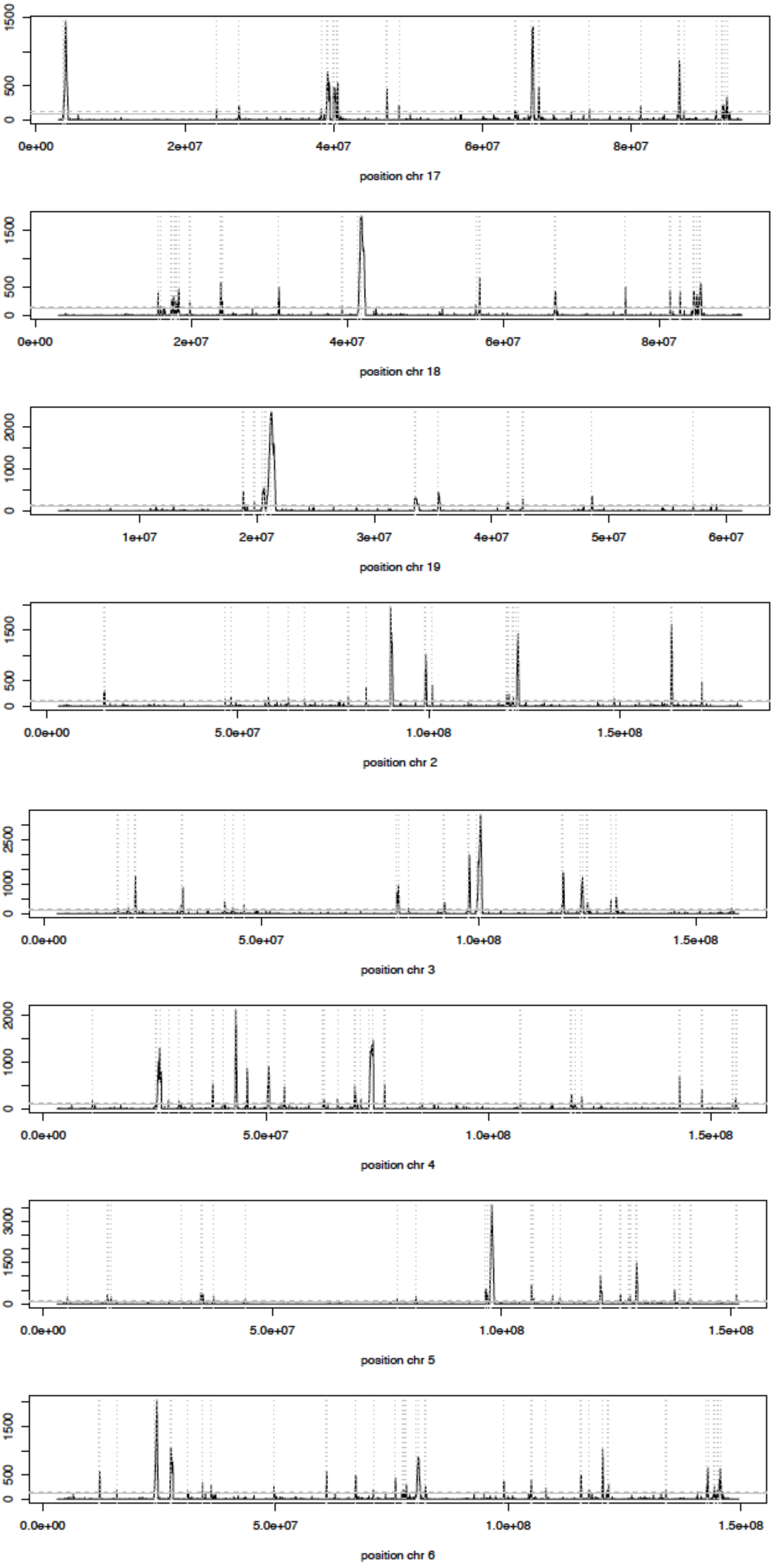

0
500
1500
0e+00
2e+07
4e+07
6e+07
8e+07
position chr 17
0
500
1500
0e+00
2e+07
4e+07
6e+07
8e+07
position chr 18
0
1000
2000
1e+07
2e+07
3e+07
4e+07
5e+07
6e+07
position chr 19
0
500
1500
0.0e+00
5.0e+07
1.0e+08
1.5e+08
position chr 2
0
1000
2500
0.0e+00
5.0e+07
1.0e+08
1.5e+08
position chr 3
0
1000
2000
0.0e+00
5.0e+07
1.0e+08
1.5e+08
position chr 4
0
1500
3000
0.0e+00
5.0e+07
1.0e+08
1.5e+08
position chr 5
0
500
1500
0.0e+00
5.0e+07
1.0e+08
1.5e+08
position chr 6

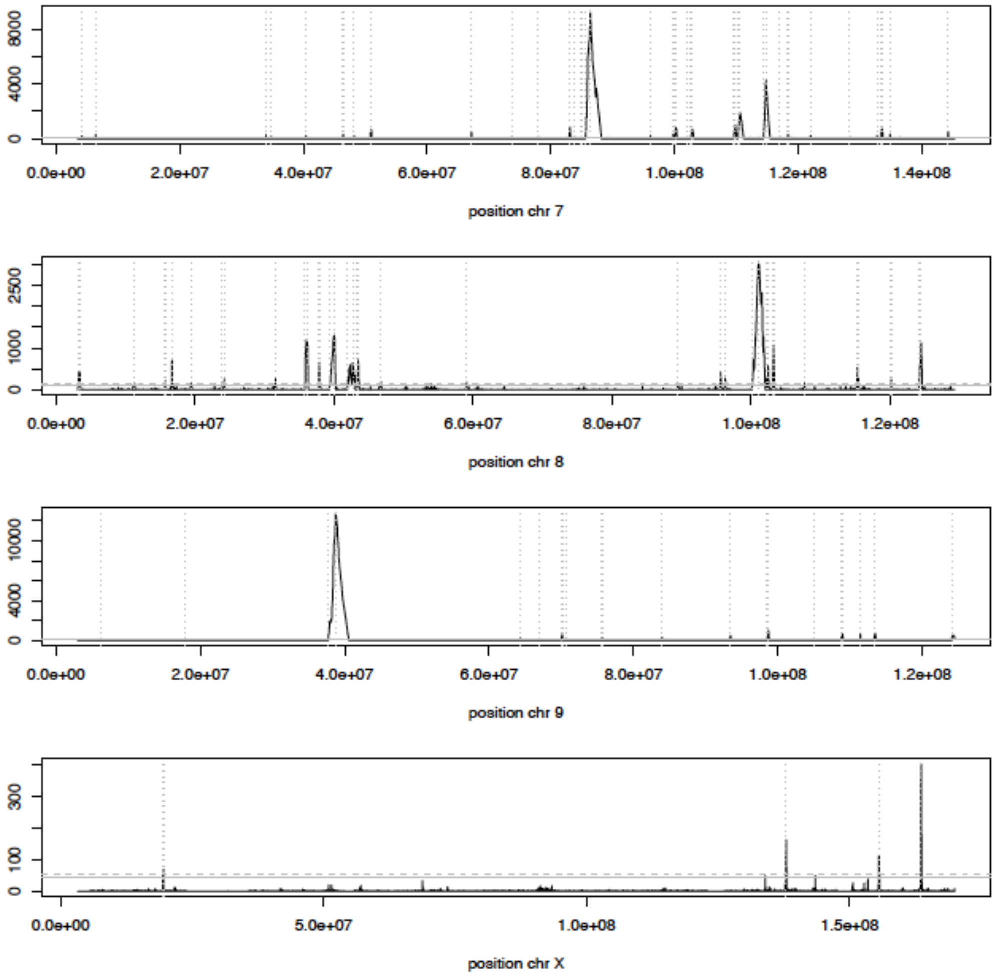


**Supplementary Figure S5**: Local score distribution along the autosomes and the X chromosome. The local score is computed as the Lindley process based on the score function $-\log_{10}(p_{FLK}) - 1$ (Fariello et al 2017). Positions are indicated in Mb. Grey plain and dotted horizontal lines indicate respectively the thresholds corresponding to the top 5% and the top 1% local score measures, computed for each chromosome. Grey dotted vertical lines indicate the centre of the region detected by the local score approach.

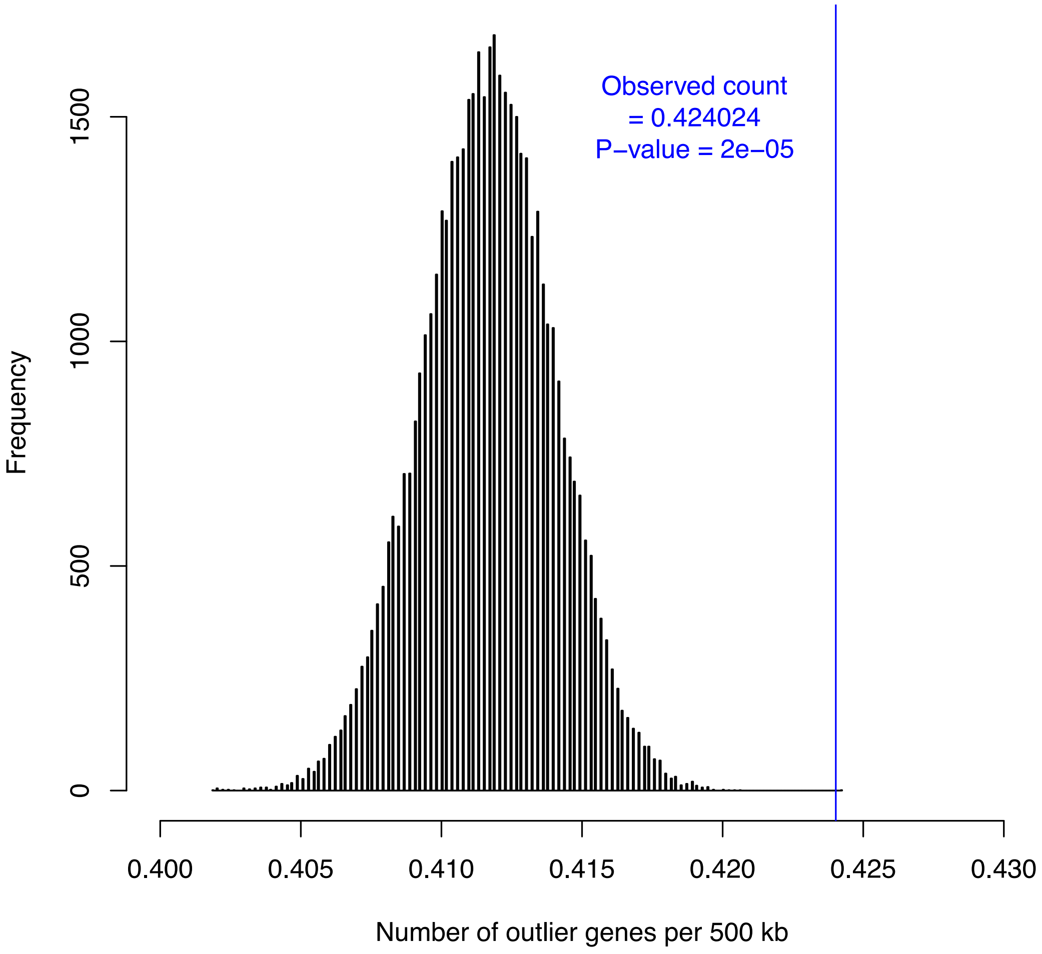


**Supplementary Figure S6** : Results of a parameter-free test for homogeneous distribution of outlier genes in the genome. Bars indicate the permutation null distribution; the blue vertical line indicates the observed average number of outlier genes per 500 kb windows. 50,000 permutations were conducted.

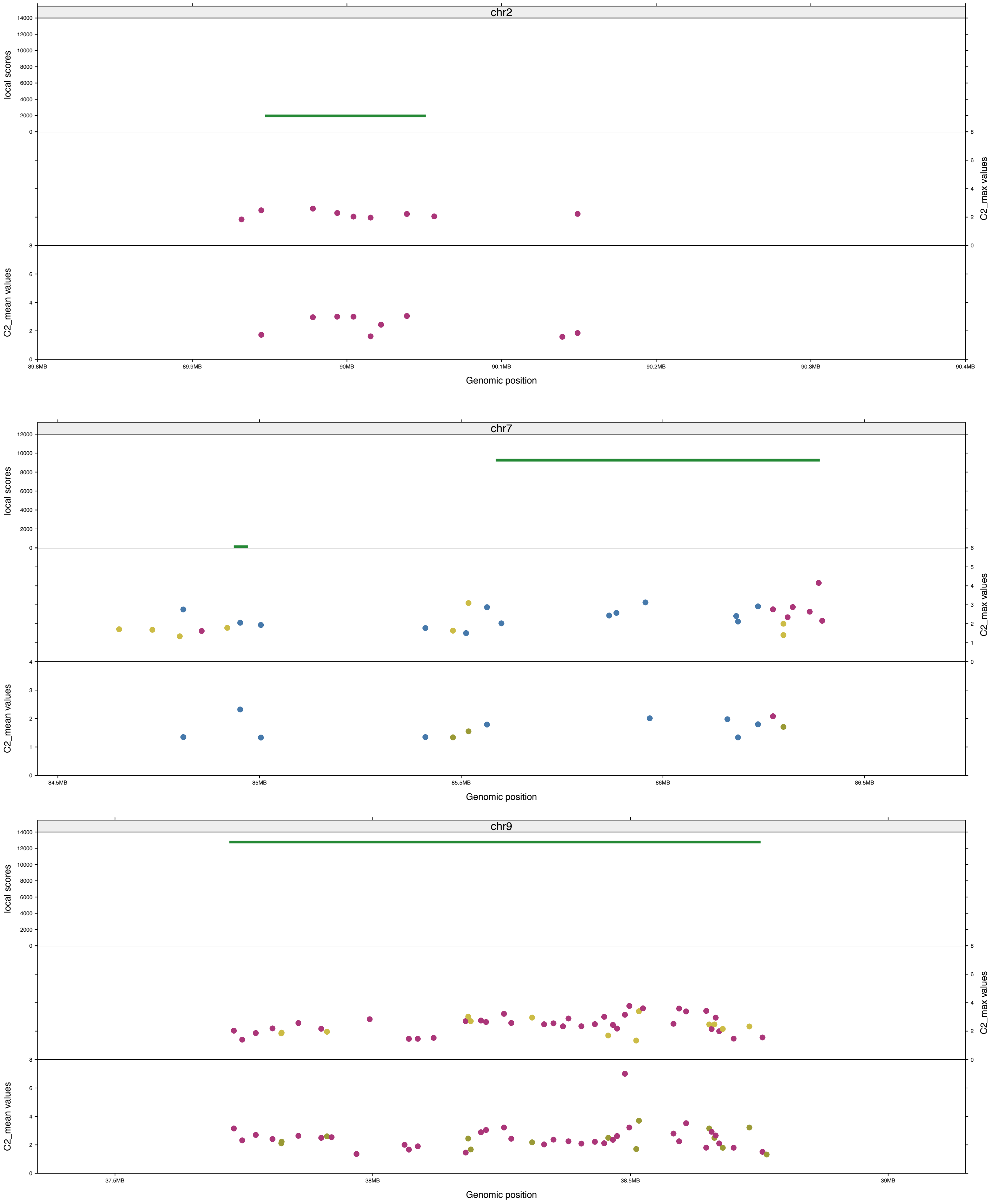


**Supplementary Figure S7**: Details of Figure 3 for the three main gene clusters on chromosomes 2, 7 and 9. The three shown tracks are (top) position and local scores for significant local score regions (green lines), (middle) position and $C_2$*_max* values for $C_2$*_max* outlier genes and (bottom) position and $C_2$*_mean* values for $C_2$*_mean* outlier genes. For $C_2$ outlier genes, *Vmn* are in blue, *Olfr* in purple, and any other type of outlier genes in light green.
